# A unified multimodal model for generalizable zero-shot and supervised protein function prediction

**DOI:** 10.1101/2025.05.09.653226

**Authors:** Frimpong Boadu, Yanli Wang, Jianlin Cheng

## Abstract

Predicting protein function is a fundamental yet challenging task that requires integrating diverse biological data modalities to capture complex functional relationships. Traditional machine learning methods often rely on single modalities or combine only a limited number (typically two), without aligning them in a unified representation, thereby constraining predictive accuracy. Moreover, most existing approaches are limited to preselected subsets of Gene Ontology (GO) function terms with sufficient annotations, making the prediction of novel function terms a persistent challenge. Here, we present FunBind, a multimodal AI model that jointly learns from protein sequences, textual descriptions, domain annotations, structural features, and GO terms to enhance prediction accuracy and infer previously unseen functions. FunBind operates in two modes: (1) self-supervised pretraining using contrastive learning to align the sequence modality with other heterogeneous modalities in a unified latent space, enabling unsupervised zero-shot function prediction, and (2) supervised fine-tuning that leverages all modalities for comprehensive and accurate function classification. Our results show that FunBind’s zero-shot capabilities allow it to generalize effectively to novel function terms never encountered before, while its joint fine-tuning strategy substantially outperforms single-modality models and current state-of-the-art approaches in prediction accuracy.

## 1 Introduction

Accurate prediction of protein function is crucial for deepening our understanding of biological processes and for advancing protein research and technological development. Most traditional approaches[1–4] rely on single-modality input data, such as protein sequences. However, single-modality methods[2, 4, 5] are unable to leverage functional information available across multiple complementary data modalities, limiting their ability to accurately predict the functions of many proteins.

There are substantial efforts to integrate a few modalities (mostly two)[1, 6, 7], such as protein sequences and structures, to improve function prediction. However, these methods simply combine the features extracted from different modalities together without semantically aligning (or binding) them in a unified representation, leading to suboptimal performance. Moreover, these methods often require all the modalities to be available to make prediction, and therefore cannot be used when some modality is missing, severely limiting their applicability.

Beyond the challenges of data integration, a significant obstacle in protein function prediction is the accurate identification of novel or poorly characterized function terms (for example, Gene Ontology (GO) terms[8, 9]) that are absent or rarely represented in existing protein function datasets and databases. The prevalent use of supervised learning methods in the field, which depend on large, labeled datasets, limits the ability to predict novel function terms that have not been encountered during training and results in poor performance on rare function terms.

To address this challenge, few-shot and zeroshot learning have emerged as promising solutions. Methods such as Tale [5] and TransFew [2] embed protein sequences and GO relationship information into a shared latent space, enabling the transfer of functional annotations from frequently annotated terms to rarely annotated ones. Similarly, DeepGOZero [10] and DeepGO-SE [7] leverage logic axioms derived from the relationships between GO terms to enhance rare or novel protein function prediction. Additionally, ProTranslator [11] integrates protein sequence data, protein-protein interactions, and textual descriptions to enhance predictions for rare function terms.

While these approaches improve the annotation of rare GO terms, they still have limitations. Many of them rely on fixed vector representations of protein function terms, which restrict their ability to generalize beyond predefined functional terms. Therefore, they still cannot accurately predict novel GO terms never seen during training. Moreover, their dependence on supervised training introduces biases linked to the quality and quantity of available labeled data, ultimately constraining their applicability to poorly annotated function terms.

To address these challenges, we introduce FunBind, a novel multimodal AI model to integrate protein sequences, structures, textual descriptions, and domain annotations for robust protein function prediction. Unlike previous approaches, FunBind effectively operates in scenarios with missing modalities by leveraging protein sequences as the central modality for cross-modality alignment, ensuring a stable foundation for learning.

Given their near-universal availability, sequences provide a reliable anchor for integrating any additional data sources, even when certain modalities are missing.

Furthermore, FunBind incorporates selfsupervised pretraining via contrastive learning to enable unsupervised zero-shot function prediction, allowing it to generalize to unseen GO terms without explicit training examples. This capability is crucial for annotating novel or poorly characterized proteins, where traditional supervised methods often fail. By learning a unified joint embedding space, FunBind captures the inherent relationships between modalities, enabling effective extrapolation to previously unseen functional terms.

We conduct extensive experiments, including contrastive learning-based multimodal alignment (binding), self-supervised pretraining, zero-shot function prediction, and supervised fine-tuning for function classification to rigorously assess FunBind’s performance. Our results demonstrate that FunBind outperforms current state-of-the-art approaches in protein function classification by effectively leveraging diverse, complementary data modalities, even when some modalities are absent. Moreover, FunBind can predict novel function terms that were never seen during training, highlighting its ability to make biologically meaningful zero-shot predictions.

## Results

### Overview of FunBind

Inspired by the vision-language model (e.g., CLIP) of aligning images and texts [12–14], we first develop a multi-modal foundational model (FunBind) to align five modalities of proteins, including sequences, structures, domain annotations in the InterPro database [15], text description, and GO function terms in the latent space by maximizing the similarity between the embedding of modalities of the same proteins and minimizing the similarity of different proteins (**Figure 1**). After the model is pre-trained by self-supervised contrastive learning, it can be directly used to predict the probability of any function term for a protein by evaluating how closely the embedding of the term is aligned with any other modalities of the protein (e.g., sequence), enabling the unsupervised zero-shot prediction of novel function terms. Moreover, FunBind is further fine-tuned via multi-label supervised learning to integrate the embeddings of all the available non-function modalities to predict function for proteins. Both the self-supervised contrastive learning and the multi-label supervised fine-tuning can work with any number of modalities, even when some modalities are missing.

**Fig. 1:**
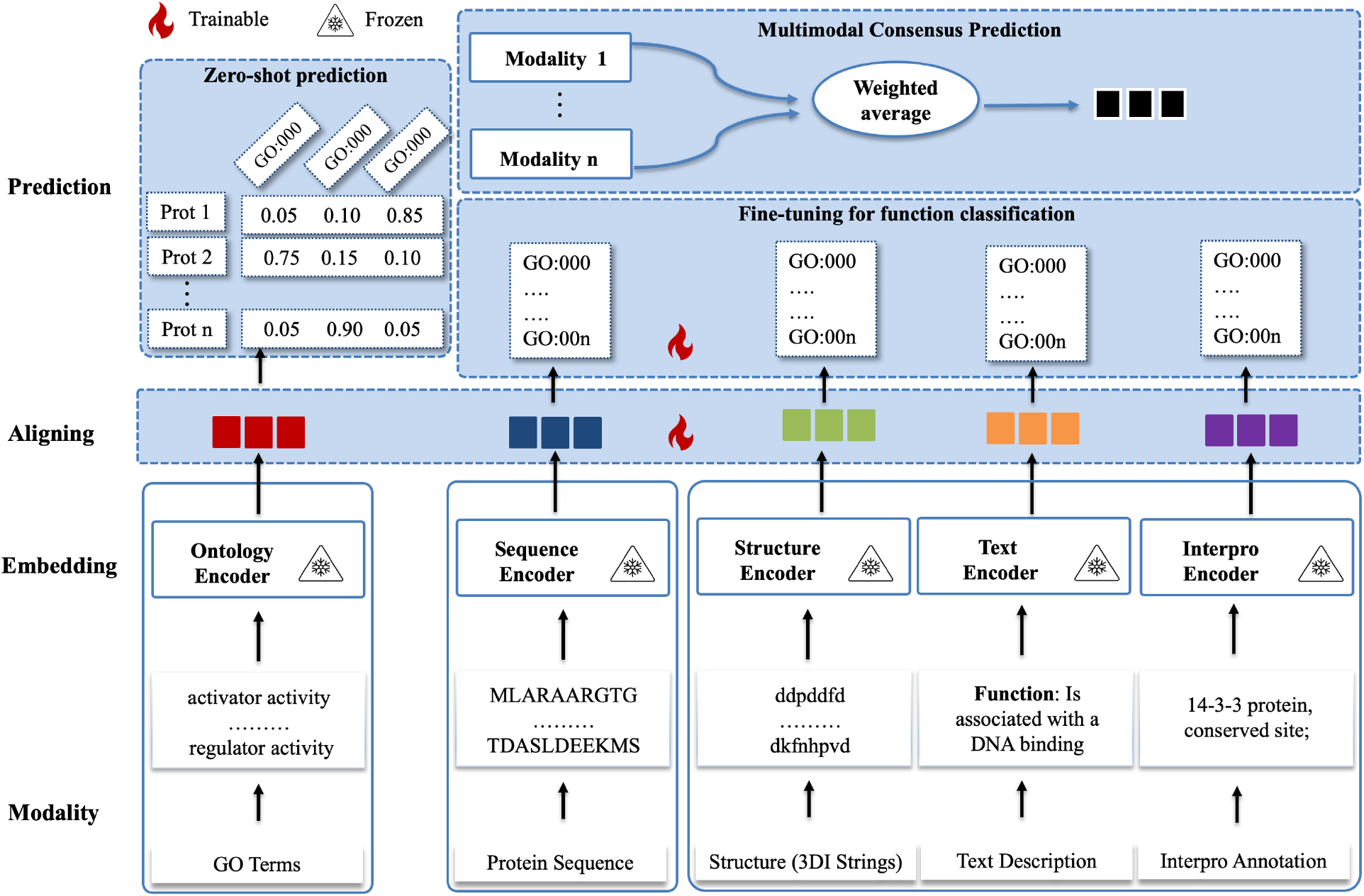
The architecture of FunBind. It integrates five data modalities — protein sequences, structures represented as 3Di strings of FoldSeek[16], textual description, and InterPro domain annotations, GO function terms via self-supervised contrastive learning. Each modality is first processed through a specialized frozen encoder to generate an embedding. The embeddings are aligned in the latent space during pretraining. After the pretraining, it can be directly used to make zero-shot prediction of protein function by evaluating the embedding of each GO term against the sequence modality (or other non-function modality) of a protein. Moreover, the aligned embedding of each non-function modality of proteins generated by the pretrained foundational model can be used by a multi-label classification head to predict their function through joint supervised fine-tuning. The predictions from the non-function modalities are then fused by weighted averaging to generate the multimodal consensus function prediction.

The objective of pretraining is to learn a unified embedding space for the five modalities, with the protein sequence modality serving as the central modality for cross-modality alignment. This is because the protein sequence is always available, while other modalities may be missing. By aligning (binding) all other four modalities with the sequence modality, the model can leverage the sequence information to learn relationships between different modalities. This approach contrasts with traditional methods that simply combine modalities without leveraging such relationships. Additionally, this strategy simplifies the learning process by requiring only *n−* 1 modality alignments rather than *n ×* (*n−* 1) pairwise alignments (n: number of modalities and n = 5 here).

FunBind’s architecture (**Figure 1**), comprises three core modules: (a) Modality encoding module to process input data from different modalities to generate embeddings, (b) Modality alignment (binding) module, to align embeddings across modalities in the latent space, and (c) Prediction module, to output functional annotations based on learned relationships, which has two submodules: (1) unsupervised zero-shot prediction and (2) supervised multi-label classification-based prediction.

For the zero-shot function prediction, the pretrained model predicts GO terms solely from alignments between a non-function modality and GO term modality (e.g., sequence-GO alignments), i.e., the GO terms that align best with the sequence modality or other modalities are selected as predicted function terms. This unsupervised approach can predict any GO function terms including novel ones never encountered before.

For the multi-label classification-based function prediction, the pretrained model is further fine-tuned by supervised learning to use available non-function modalities to predict function.

Here, the aligned embedding of each available nonfunction modality is used by a multi-label classification head to predict protein functions within a fixed set of pre-determined GO terms. The predictions from all available non-function modalities are averaged to generate the multimodal consensus prediction.

### Zero-Shot Prediction of Protein Function Using Sequence Modality

FunBind employs a zero-shot inference strategy to predict protein function, inspired by multi-modal vision-language models [12–14]. The pretrained FunBind first generates aligned embeddings for protein sequences and any GO terms under consideration in a shared latent embedding space, then computes cosine similarity between the embeddings, and finally assigns the GO terms with the highest similarity scores as predicted functions for the proteins.

To evaluate zero-shot performance, we curated a test set (Test Zero) (see **Table 1** for its statistics and the “Methods” Section for details), consisting of new proteins, each annotated with at least one novel GO term absent in the training data. The statistics shows that all the proteins have sequence modality, but only a portion of them have other non-sequence and non-function modalities in this dataset. To fairly compare the zero-shot prediction performance of different modalities, we focus on describing the results of the zero-shot prediction on the proteins having all modalities available (as represented in the Intersection column in **Table 1**), while referring readers to the results on all the proteins in Test Zero in the supplementary materials.

**Table 1:** The statistics of Test Zero Dataset for zero-shot prediction. The number of proteins having each of four modalities for each function ontology / category (CC: cellular component, MF: molecular function, and BP: biological process) is reported. The Intersection column lists the number of proteins having all four non-function modalities. The All row lists the sum of the numbers of proteins in CC, MF and BP combined.

| Ontology | Sequence | Structure | Text | InterPro | Intersection |
| --- | --- | --- | --- | --- | --- |
| CC | 73 | 59 | 73 | 70 | 56 |
| MF | 192 | 43 | 49 | 50 | 42 |
| BP | 427 | 92 | 109 | 105 | 84 |
| All | 659 | 185 | 215 | 209 | 173 |

The test proteins were randomly divided into groups (batches), each containing 10–12 proteins and the same number of novel GO terms. Each protein was associated with a single novel GO term, which was used as its label. Within each group, all the protein sequences were aligned with all candidate GO terms to calculate the similarity between each protein sequence and each GO term, which is used to rank GO terms for each protein. This setup is similar to how the zero-shot prediction was evaluated with vision-language models [12]. To account for variability in random grouping, the experiment was repeated 10 times. In each trial, proteins were randomly grouped, ensuring that the results reflect the model’s performance across different groupings and reducing the potential bias from random selection. The mean and standard deviation of performance metrics are reported for each function category as well as for the All function categories (**Table 2**). The performance was evaluated using Recall@k (R@1, R@3, R@5), which measures the fraction of proteins with correctly retrieved novel GO terms among the top-k predictions, and Mean Reciprocal Rank (MRR), which considers the ranking position of relevant GO terms.

**Table 2:** The performance of zero-shot function prediction using sequence modality as query. The mean and standard deviation of the recall at 1, 3, and 5 predictions, and Mean Reciprocal Rank (MRR) for 10 trials on the proteins in Test Zero dataset.

| Ontology | R@1 | R@3 | R@5 | MRR |
| --- | --- | --- | --- | --- |
| CC | 0.7548<br>($\pm 0.0254$) | 0.8732<br>($\pm 0.0124$) | 0.9129<br>($\pm 0.0201$) | 0.8302<br>($\pm 0.0129$) |
| MF | 0.6357<br>($\pm 0.0564$) | 0.8405<br>($\pm 0.0302$) | 0.9048<br>($\pm 0.0213$) | 0.7550<br>($\pm 0.0312$) |
| BP | 0.4821<br>($\pm 0.0374$) | 0.6869<br>($\pm 0.0160$) | 0.7845<br>($\pm 0.0180$) | 0.6225<br>($\pm 0.0206$) |
| All | 0.5947<br>($\pm 0.0280$) | 0.7969<br>($\pm 0.0111$) | 0.8749<br>($\pm 0.0105$) | 0.7180<br>($\pm 0.0161$) |

As shown in **Table 2**, across all three function categories (CC, MF, and BP), recall improves as more predictions per protein are considered. The recall for one prediction per protein (R@1) ranges from 0.4821 for BP, 0.6357 for MF, to 0.7548 for CC, and increases to 0.7845 for BP, 0.9048 for MF, and 0.9129 for CC, when making five predictions per protein (R@5). These results demonstrate FunBind can rather accurately predict novel GO terms not encountered during training.

Among the three GO function categories, CC exhibits the highest retrieval performance across all metrics, followed by MF and then BP. This performance disparity may be partly due to the level of the clarity in the definition of the GO terms in each ontology. CC and MF terms at the same level in the gene ontology graph may be more distinct from each other than BP terms, making them easier to differentiate. For example, two different CC terms at the same level refer to two completely different cellular locations without any overlap, while some BP terms may have some semantic similarity.

To illustrate how the zero-shot function prediction selects GO terms for a protein, **Figure 2(a)** visualizes the protein sequence–GO term alignment similarity for each of three randomly selected groups of proteins for all function categories (see supplementary **Figure S1** for the alignments for the remaining 12 groups). Each heatmap in **Figure 2(a)** visualizes the similarity between the proteins and the novel GO terms in a group. The proteins and their corresponding GO terms are arranged sequentially in the same order, with each element on the diagonal of the heatmap representing the similarity score between each protein and its corresponding true GO term (label) and each element off the diagonal representing the similarity score between each protein and a GO term not associated with it. It is shown that many elements on the diagonal have relatively higher similarity scores, highlighting the strong alignment between protein sequences and their labels, which leads to a correct prediction. It is worth noting that the zero-shot prediction of novel function terms is done fully through the unsupervised contrastive learning without using label-based supervised learning employed by most machine learning methods for function prediction.

**Fig. 2:**
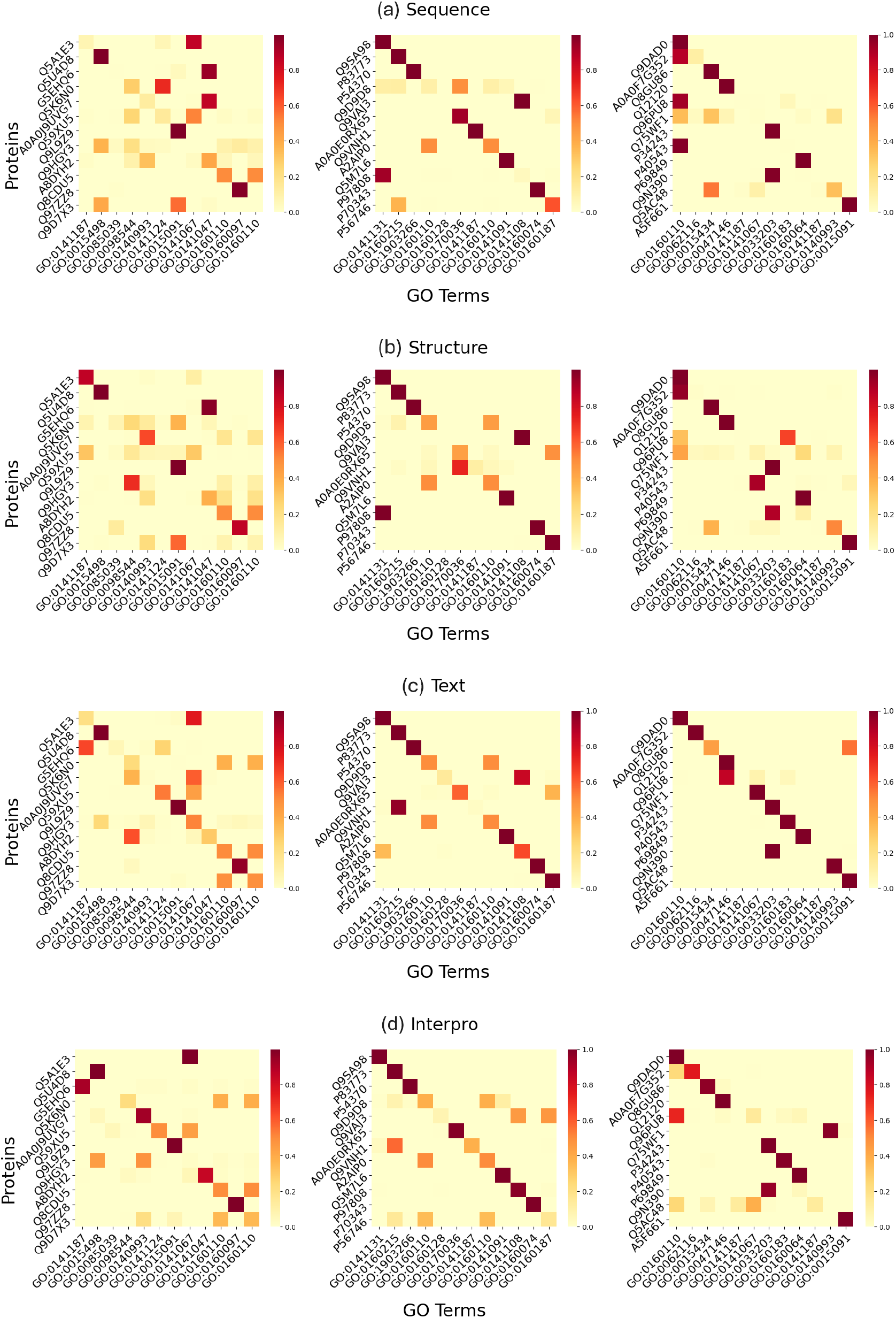
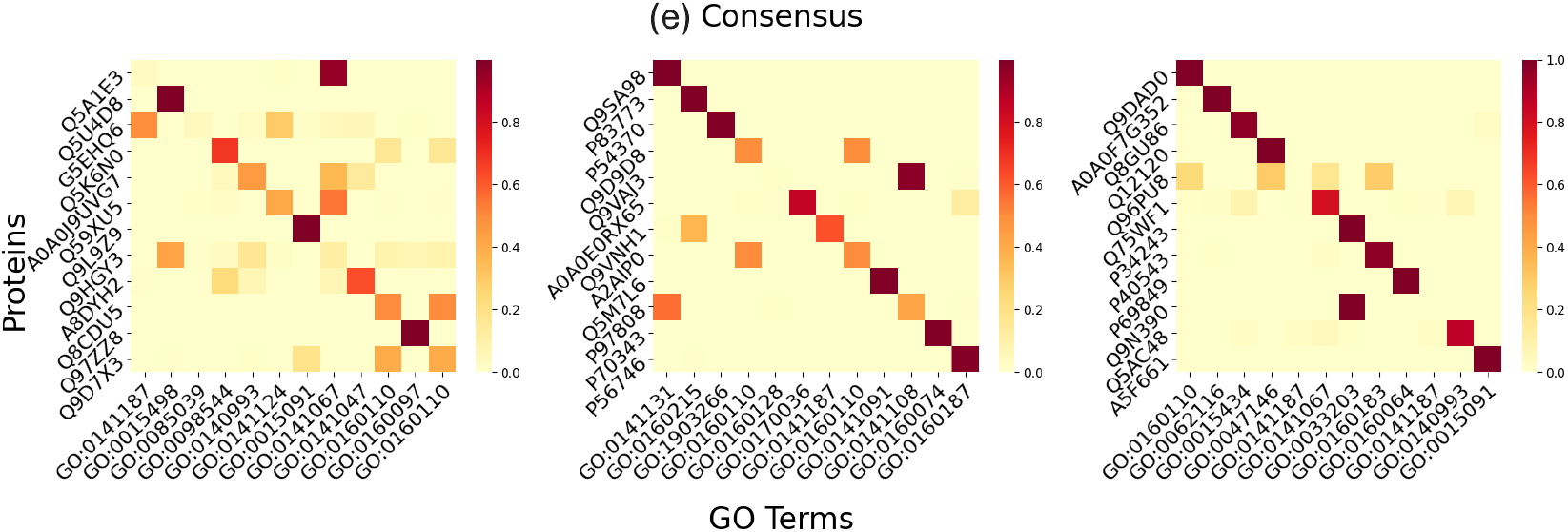
Heatmaps illustrating the alignment similarity between the embeddings of each non-function modality of the proteins (rows) and the embeddings of the novel GO terms (columns) for three randomly partitioned groups in zero-shot prediction. The GO terms are ordered in the same way as the proteins so that the *i*th GO term is the only true GO term of the *i*th protein. Therefore, the values on the diagonal reflect the similarity between a non-function modality of the proteins and their true GO terms, higher the better. **(a)** Sequence modality, **(b)** Structure modality, **(c)** Text modality, **(d)** Interpro domain annotation modality, and **(e)** Consensus of the four modalities..

### Zero-Shot Function Prediction Using Other Non-Central Modalities

During the pretraining of FunBind, protein sequence serves as the central modality to be aligned with other modalities of proteins (Textual description, Structures, InterPro domain annotations, and GO terms). There is no direct alignment between GO terms and other non-sequence modalities. However, we hypothesize that the other non-sequence modalities and GO terms have been aligned implicitly through the bridge of the central sequence modality, a phenomenon known as emergent binding in the vision-language models [13, 14].

To investigate this emergent binding, we evaluated FunBind’s zero-shot prediction of using structural data, textual descriptions, and InterPro domain annotations as queries to predict GO terms, respectively. This evaluation follows the same methodology as the sequence-based zero-shot prediction, where the cosine similarity between modality embeddings and GO term embeddings is used to rank and select GO terms for a protein. The performance of zero-shot function prediction using each non-sequence modality was tested on the same test proteins used in the sequence-based zero-shot evaluation, which have all five data modalities available.

The results of this “emergent binding” evaluation are presented in **Table 3**. Text modality consistently achieves the highest scores across all metrics and function categories. It even outperforms the central sequence modality in most cases (**Table 2**), highlighting the informativeness of textual descriptions for protein function prediction when they are available. Furthermore, text exhibits the most stable performance, with consistently low standard deviations across all function ontologies.

**Table 3:** Zero-shot prediction performance of FunBind using non-central modalities.

| Ontology | Text |  |  |  | Interpro |  |  |  | Structure |  |  |  |
| --- | --- | --- | --- | --- | --- | --- | --- | --- | --- | --- | --- | --- |
|  | R@1 | R@3 | R@5 | MRR | R@1 | R@3 | R@5 | MRR | R@1 | R@3 | R@5 | MRR |
| **CC** | 0.8239<br>($\pm 0.0184$) | 0.8874<br>($\pm 0.0081$) | 0.9000<br>($\pm 0.0081$) | 0.8697<br>($\pm 0.0093$) | 0.7521<br>($\pm 0.0245$) | 0.8283<br>($\pm 0.0092$) | 0.8552<br>($\pm 0.0142$) | 0.8127<br>($\pm 0.0129$) | 0.6694<br>($\pm 0.0274$) | 0.7839<br>($\pm 0.0093$) | 0.8179<br>($\pm 0.0150$) | 0.7524<br>($\pm 0.0184$) |
| **MF** | 0.6976<br>($\pm 0.0302$) | 0.8810<br>($\pm 0.0282$) | 0.9429<br>($\pm 0.0218$) | 0.8014<br>($\pm 0.0159$) | 0.6667<br>($\pm 0.0499$) | 0.8571<br>($\pm 0.0151$) | 0.9310<br>($\pm 0.0310$) | 0.7774<br>($\pm 0.0244$) | 0.6262<br>($\pm 0.0533$) | 0.8643<br>($\pm 0.0354$) | 0.9167<br>($\pm 0.0160$) | 0.7546<br>($\pm 0.0276$) |
| **BP** | 0.4738<br>($\pm 0.0310$) | 0.7226<br>($\pm 0.0292$) | 0.8452<br>($\pm 0.0238$) | 0.6304<br>($\pm 0.0204$) | 0.4583<br>($\pm 0.0234$) | 0.6905<br>($\pm 0.0292$) | 0.7905<br>($\pm 0.0208$) | 0.6094<br>($\pm 0.0152$) | 0.4619<br>($\pm 0.0158$) | 0.6774<br>($\pm 0.0202$) | 0.7679<br>($\pm 0.0179$) | 0.6050<br>($\pm 0.0089$) |
| **All** | 0.6538<br>($\pm 0.0232$) | 0.8216<br>($\pm 0.0124$) | 0.8874<br>($\pm 0.0081$) | 0.7575<br>($\pm 0.0124$) | 0.6297<br>($\pm 0.0222$) | 0.7891<br>($\pm 0.0157$) | 0.8608<br>($\pm 0.0120$) | 0.7341<br>($\pm 0.0134$) | 0.5714<br>($\pm 0.0249$) | 0.7571<br>($\pm 0.0147$) | 0.8420<br>($\pm 0.0121$) | 0.6924<br>($\pm 0.0142$) |

InterPro domain annotation modality (**Table 3**) performs similarly as the central sequence modality (**Table 2**), and better than the structure modality in CC and MF but similarly in BP, indicating that domain and family annotations offer valuable functional insights. Even though the zero-shot prediction performance of the structure modality is relatively weaker, it is complementary with other modalities because combining them can yield better results (see the results in the next section).

**Figure 2(b)(c)(d)** visualize the alignment similarity between each non-central modality and GO terms for three randomly selected groups respectively, showing that the modality of many proteins are aligned well with their true GO terms. The visualization of the similarity matrices for the remaining groups for the structure, text and Interpro domain annotation modalities is provided in supplementary **Figures S2-S4**.

Finally, in contrast to dividing the test proteins into smaller, ontology-associated groups in the experiments above, we also used the entire set of test proteins as one single group (batch) for FunBind to perform zero-shot prediction. The results for the subset of proteins with data available for all four non-function modalities are reported in supplementary **Figure S6** and those for the entire protein set (regardless of modality availability) are reported in supplementary **Figure S7**. The zeroshot approach still predicts the novel terms of the proteins well in these settings, even though the accuracy is lower than dividing the proteins into smaller groups. The reason is that treating all the proteins as one single group requires retrieving true GO terms from more GO term candidates.

### Consensus Zero-Shot Function Prediction by Combining Multiple Modalities

To study if combining four non-function modalities can improve zero-shot function prediction, we performed a grid search of weights to combine modality-specific similarity scores through a weighted summation. We tested different combinations of weights assigned to the modalities, with each weight ranging from 0 to 1 in steps of 0.1, while maintaining the sum of weights equals 1. Consistent with previous experiments, we used the same test proteins with all the modalities available in Test Zero dataset and the same experimental protocol to test the consensus zero-shot approach. The configuration that achieves the highest Recall@1 score is selected for combining the modalities in the final evaluation. The optimal weights for the four non-function modalities are reported in supplementary **Table S2**, with the corresponding zero-shot predictioin results shown in **Table 4**. It is shown that the consensus zero-shot prediction outperforms each individual modality (**Table 3**).

**Table 4:** Mean and standard deviation of Recall@1, Recall@3, Recall@5, and Mean Reciprocal Rank (MRR) for consensus zero-shot function prediction by combining sequence, structure, text, and InterPro modalities. Results are averaged over 10 trials on the Test Zero dataset.

| Ontology | R@1 | R@3 | R@5 | MRR |
| --- | --- | --- | --- | --- |
| **CC** | 0.8633<br>($\pm 0.0172$) | 0.9252<br>($\pm 0.0077$) | 0.9321<br>($\pm 0.0106$) | 0.9019<br>($\pm 0.0100$) |
| **MF** | 0.7452<br>($\pm 0.0465$) | 0.9262<br>($\pm 0.0310$) | 0.9738<br>($\pm 0.0071$) | 0.8377<br>($\pm 0.0268$) |
| **BP** | 0.5333<br>($\pm 0.0327$) | 0.7286<br>($\pm 0.0128$) | 0.8060<br>($\pm 0.0169$) | 0.6631<br>($\pm 0.0167$) |
| **All** | 0.6939<br>($\pm 0.0166$) | 0.8293<br>($\pm 0.0077$) | 0.8922<br>($\pm 0.0098$) | 0.7838<br>($\pm 0.0101$) |

**Figure 2(e)** visualizes the alignment similarity of the consensus approach between the proteins and the ground-truth GO terms in three randomly selected groups. Overall, integrating predictions from all four modalities yields stronger alignment with the ground truth than using individual modalities alone. Similarity matrices for the remaining groups are provided in Supplementary **Figure S5**.

### Case Study of Zero-Shot Function Prediction

We use four examples to illustrate FunBind’s zeroshot capability of predicting protein function. We selected four proteins (UniProt ID: **A8BPK8, P18335, Q12198**, and **Q64565**) along with their 12 Gene Ontology (GO) terms from the Text Zero dataset. One protein may have one or a few true GO terms among the 12 terms.

For each protein and each available modality, FunBind encoded both the modality and the corresponding GO terms, and subsequently computed the cosine similarity between the modality embedding and each GO term embedding. These similarity scores were transformed into confidence scores by a softmax function, which were then multiplied by 100 as percentages.

**Figure 3** shows the results. Each bar chart displays the predicted GO terms for a protein, with the height of each bar representing FunBind’s confidence in that prediction.

**Fig. 3:**
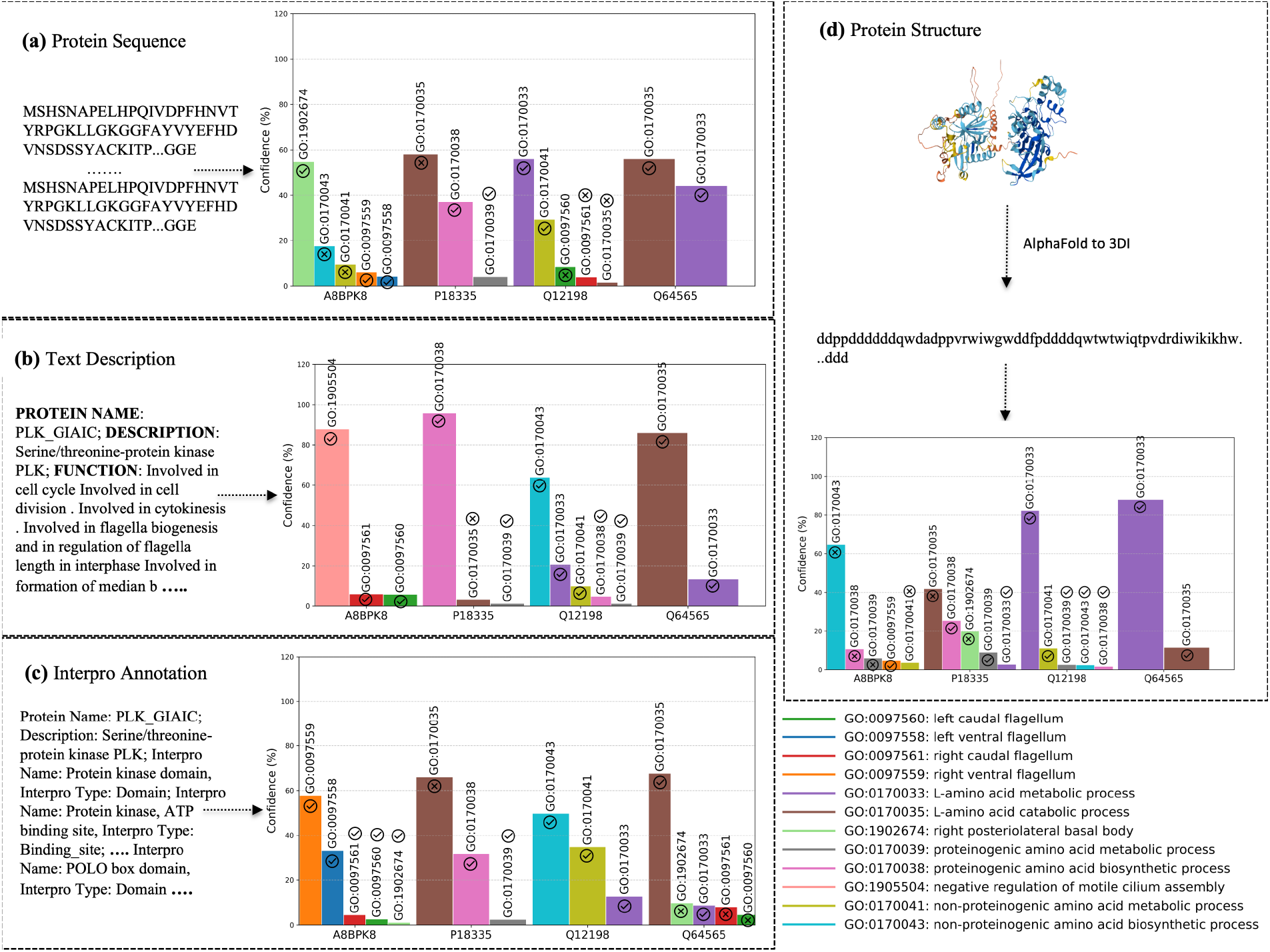
The zero-shot function predictions for four protein examples made by the four modalities of FunBind respectively. **(a)** protein sequence. **(b)** textual description. **(c)** InterPro domain annotation. **(d)** protein structure. The color-coded legends denote 12 different function terms. Two markers shown on top of each bar indicate if a predicted GO term is correct or not.

The Text modality made correct top-1 prediction for all the four proteins, while the Sequence and Interpro modalities retrieved the first GO term correctly for three out of four proteins and the Structure modality made correct first GO term retrieval for two proteins, highlighting the model’s ability to prediction function across modalities.

Certain biological functions are consistently retrieved across multiple modalities. For instance, L-amino acid metabolic process (GO:0170033) and L-amino acid catabolic process (GO:0170035) are correctly retrieved within top five predictions by all four modalities for protein Q64565, showing the consistency of these functional predictions.

However, certain function terms are only retrieved by specific modalities, indicating modality biases and complementarity. For instance, for protein A8BPK8, each of three modalities (Text, Sequence, and Interpro domain) retrieved one different correct GO term as top-1 prediction, while the Structure modality failed to select the first GO term correctly.

### Comparison of Fine-tuned FunBind and Existing Methods on Test All Dataset

As shown in (**Figure 1**), we also fine-tuned the pretrained FunBind on the training dataset via supervised learning to predict protein function and compared it with five existing methods on the Test All dataset. Test All comprises the new proteins released after the latest release date of the proteins in the training data (see “Methods” section for details). The statistics of Test All is shown in **Table 5**.

**Table 5:** Statistics of Test All Dataset.

| Ontology | Sequence | Structure | Text | InterPro |
| --- | --- | --- | --- | --- |
| CC | 1883 | 1728 | 1828 | 1818 |
| MF | 2441 | 2217 | 2391 | 2355 |
| BP | 2585 | 2347 | 2522 | 2465 |

Specifically, the fine-tuned FunBind was compared with two traditional baseline methods (Naive, DiamondBlast[3]), three current state-of-the-art (SOTA) deep learning methods (TransFew[2], SPROF-GO[6] and DeepGO-SE [7]), the predictions of the four non-function modalities, in terms of multiple metrics, including F_*max*_, Area under the Precision-Recall curve (AUPR), weighted F_*max*_, and S_*min*_ of measuring the uncertain/missing information in function predictions [17–21] (see the detailed definition of the evaluation metrics and a summary of the five function prediction methods in supplementary **Notes 1 and 2** respectively). FunBind uses the multimodal consensus of all four non-function modalities (Sequence, Text, InterPro, and Structure) as prediction. The weights for the predictions of the modalities in FundBind were optimized on the validation dataset before being blindly used on the test dataset.

For SPROF-GO, we obtained predictions from its webpage (as of February 9, 2025), while for DeepGOSE, TransFew, Naive and DiamondBlast, we generated their predictions by running them locally. FundBind, TransFew, SPROF-GO, DeepGO-SE, Naive, and the Sequence modality made predictions for all the proteins (coverage = 1), while the other individual modalities made predictions for 90% to 97% of proteins because some modalities are missing for a small portion of proteins. DiamondBlast only made predictions for 75-78% of proteins in three function categories because it could not found homologous hits for some proteins.

The results of FunBind, the individual modalities of FunBind, and the five existing methods on the Test All dataset are presented in **Table 6**. The Precision-Recall curves of the methods are shown in **Figure 4**. The results show that FunBind outperforms all the existing methods and four individual modalities, in terms of all the metrics for all three function categories (CC, MF, and BP). It is clearly demonstrated that integrating multiple modalities with FundBind advances the state of the art of protein function prediction. FundBind not only consistently outperform three state-of-the-art deep learning methods (TransFew[2], SPROF-GO[6] and DeepGO-SE [7]) across the board, but also improve the prediction accuracy by a large margin in some situations. For instance, the AUPR of FundBind for BP is 0.429, more than 10 percentage points higher than 0.320, 0.329, 0.283 of DeepGO-SE, SPROF-GO and TransFew.

**Table 6:** Performance of FunBind and existing methods on Test All dataset. Bold font highlights the best performance and underline denotes the second best performance.

| Methods | | $F_{max}$ (↑) | | | $WF_{max}$ (↑) | | | $AUPR$ (↑) | | | $S_{min}$ (↓) | | | Coverage | | |
| --- | --- | --- | --- | --- | --- | --- | --- | --- | --- | --- | --- | --- | --- | --- | --- | --- |
|  |  | CC | MF | BP | CC | MF | BP | CC | MF | BP | CC | MF | BP | CC | MF | BP |
| Baseline | Naive | 0.588 | 0.534 | 0.267 | 0.323 | 0.372 | 0.183 | 0.483 | 0.282 | 0.144 | 6.351 | 5.034 | 17.532 | 1.0 | 1.0 | 1.0 |
|  | DiamondBLAST | 0.587 | 0.563 | 0.347 | 0.470 | 0.480 | 0.292 | 0.050 | 0.047 | 0.043 | 7.566 | 5.146 | 23.342 | 0.78 | 0.76 | 0.75 |
| Current SOTA | DeepGO-SE | 0.681 | 0.634 | 0.402 | 0.502 | 0.519 | 0.323 | 0.667 | 0.552 | 0.320 | 5.481 | 4.020 | 16.085 | 1.0 | 1.0 | 1.0 |
|  | SPROF-GO | 0.718 | 0.680 | 0.414 | 0.579 | 0.577 | 0.334 | <u>0.747</u> | 0.619 | 0.329 | 4.922 | 3.518 | 15.476 | 1.0 | 1.0 | 1.0 |
|  | TransFew | 0.703 | 0.680 | 0.421 | 0.570 | 0.593 | 0.349 | 0.631 | 0.528 | 0.283 | 5.182 | 3.735 | 16.173 | 1.0 | 1.0 | 1.0 |
| Single modal | Structure | 0.671 | 0.648 | 0.413 | 0.526 | 0.563 | 0.335 | 0.659 | 0.546 | 0.299 | 5.352 | 3.801 | 15.660 | 0.91 | 0.90 | 0.90 |
|  | Interpro | 0.697 | 0.686 | 0.457 | 0.560 | 0.604 | 0.386 | 0.714 | 0.603 | 0.355 | 5.084 | 3.507 | 14.747 | 0.96 | 0.96 | 0.95 |
|  | Sequence | 0.695 | 0.663 | 0.449 | 0.557 | 0.570 | 0.375 | 0.516 | 0.491 | 0.330 | 5.413 | 3.853 | 15.472 | 1.0 | 1.0 | 1.0 |
|  | Text | <u>0.721</u> | <u>0.697</u> | <u>0.495</u> | <u>0.601</u> | <u>0.609</u> | <u>0.431</u> | 0.736 | <u>0.626</u> | <u>0.407</u> | <u>4.589</u> | <u>3.444</u> | <u>13.889</u> | 0.97 | 0.98 | 0.97 |
|  | FunBind | **0.738** | **0.714** | **0.505** | **0.618** | **0.631** | **0.437** | **0.776** | **0.665** | **0.429** | **4.485** | **3.343** | **13.758** | 1.0 | 1.0 | 1.0 |

**Note:** FunBind: multimodal consensus with sequence, structure, text and interpro.

**Fig. 4:**
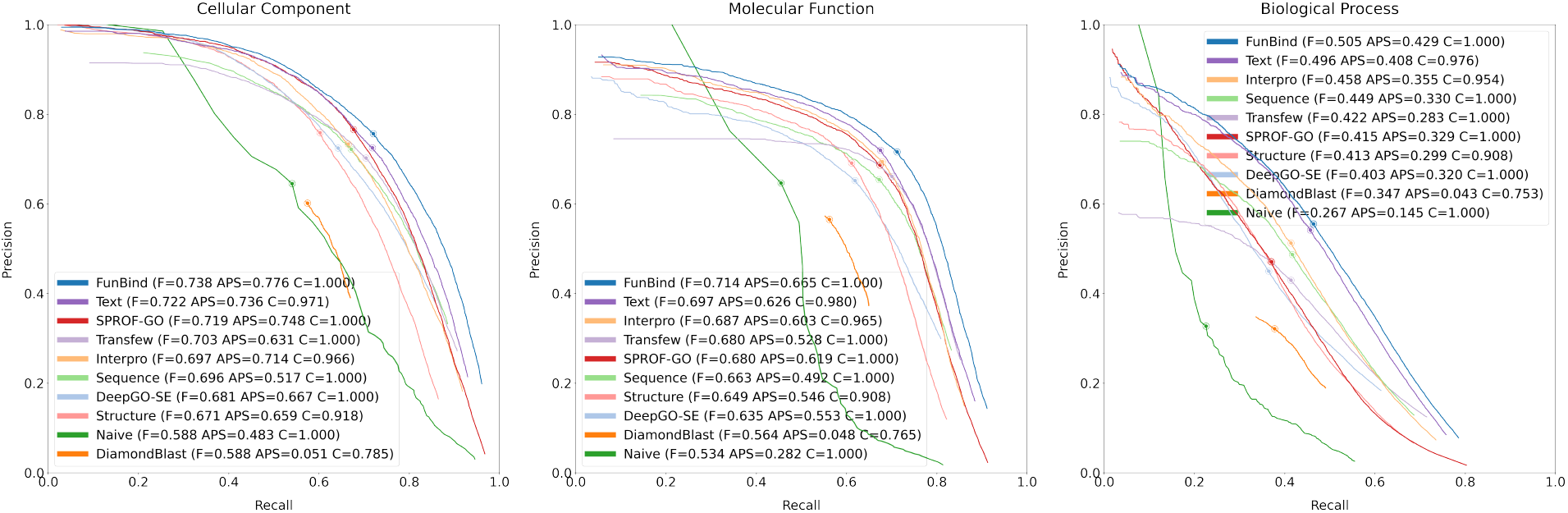
The Precision-Recall curves of FunBind and existing methods for the three ontologies (BP, MF, and CC) on the Test All dataset. The circled dot marks the point where each method achieves the highest *F*_*max*_. Each plot also includes the AUPR score and the coverage of each method (the percent of proteins which a method can make prediction for). FunBind outperforms all existing methods, particularly leading in BP by a large margin.

Among the four individual modalities, on their own set of proteins on which they were tested, Text performs best, followed by InterPro domain annotations, Sequence, and Structure. Text excels in capturing functional information, likely benefiting from rich descriptive annotations. All the four individual modalities have substantially higher scores than the two baseline methods (Navie and DiamondBLAST). The scores of Text, Sequence, and Interpro are also on par with the three deep learning methods. Particularly, Text consistently has higher scores than them in terms of almost all the metrics. However, the comparison between the four individual modalities and the other methods here is not fully fair because some of them did not make predictions for all the proteins (i.e., coverage *<* 1).

For BP, among the four individual modalities, Text achieves the highest performance. Integrating it with other modalities yields only a slight improvement, indicating that textual annotations are the primary contributors to the significant accuracy gains in BP prediction in FunBind. Meanwhile, Sequence exhibits the highest coverage, supporting its role as the central modality for integrating the others.

### Performance of Fine-tuned FunBind and Existing Methods on Novel Test Proteins

To assess how well FunBind generalizes to novel proteins that are very dissimilar to the training proteins, we compare FundBind and the existing methods on the Text Novel dataset (see its statistics in **Table 7**), which is a subset of Test All that contain only proteins that have less than 30% sequence identity with any protein in the training data. The results of all the methods are shown in **Table 8**. The Precision-Recall curves of the methods are shown in **Figure 5**.

**Table 7:** Statistics of Test Novel Dataset.

| Ontology | Sequence | Structure | Text | InterPro |
| --- | --- | --- | --- | --- |
| CC | 206 | 169 | 179 | 166 |
| MF | 231 | 179 | 210 | 199 |
| BP | 282 | 213 | 256 | 225 |

**Table 8:**
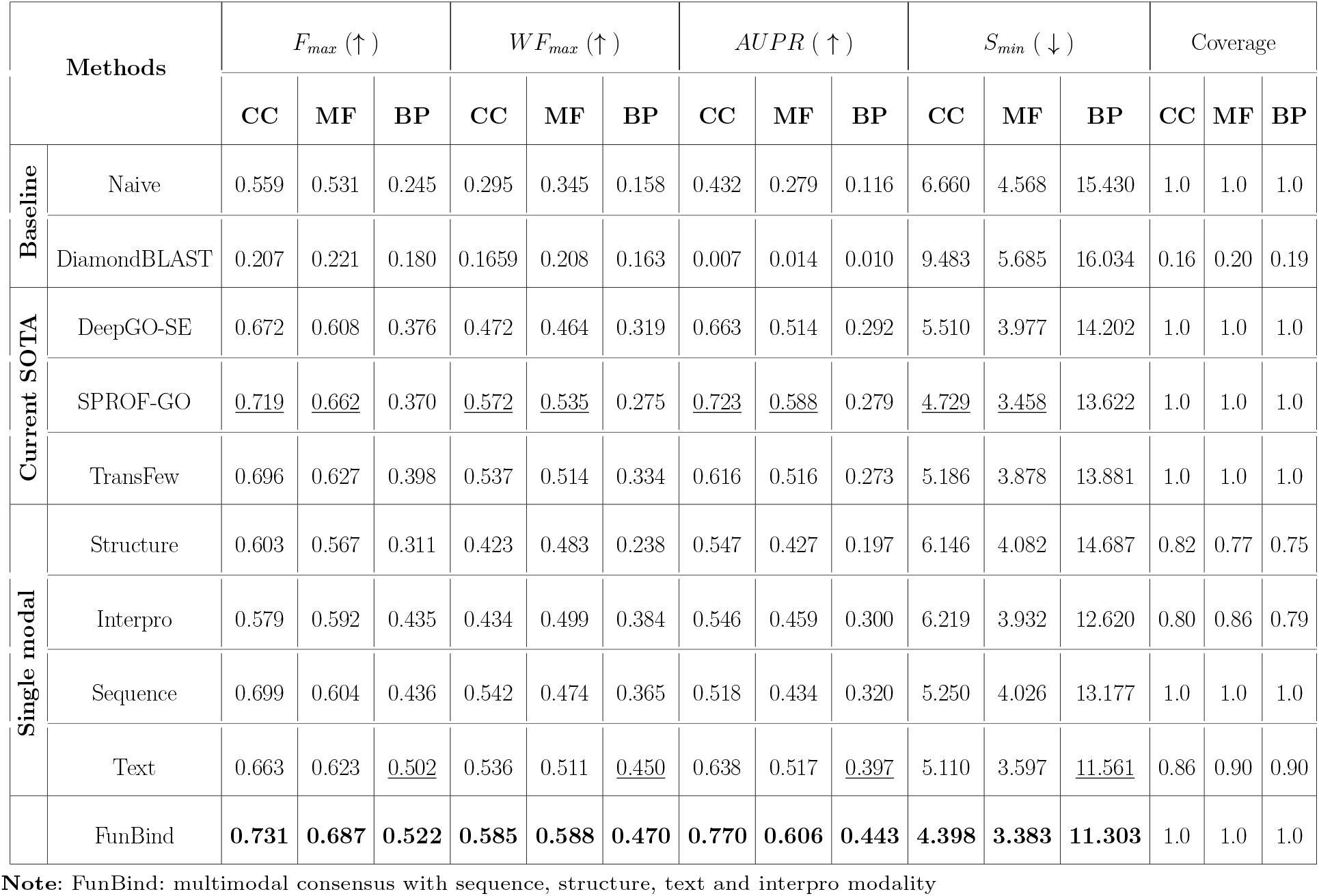
Performance of FunBind and existing methods on Test Novel Dataset. Bold font highlights the best performance and underline denotes the second best performance.

| Methods | | $F_{max} (\uparrow)$ | | | $WF_{max} (\uparrow)$ | | | $AUPR (\uparrow)$ | | | $S_{min} (\downarrow)$ | | | Coverage | | |
| --- | --- | --- | --- | --- | --- | --- | --- | --- | --- | --- | --- | --- | --- | --- | --- | --- |
|  |  | CC | MF | BP | CC | MF | BP | CC | MF | BP | CC | MF | BP | CC | MF | BP |
| Baseline | Naive | 0.559 | 0.531 | 0.245 | 0.295 | 0.345 | 0.158 | 0.432 | 0.279 | 0.116 | 6.660 | 4.568 | 15.430 | 1.0 | 1.0 | 1.0 |
|  | DiamondBLAST | 0.207 | 0.221 | 0.180 | 0.1659 | 0.208 | 0.163 | 0.007 | 0.014 | 0.010 | 9.483 | 5.685 | 16.034 | 0.16 | 0.20 | 0.19 |
| Current SOTA | DeepGO-SE | 0.672 | 0.608 | 0.376 | 0.472 | 0.464 | 0.319 | 0.663 | 0.514 | 0.292 | 5.510 | 3.977 | 14.202 | 1.0 | 1.0 | 1.0 |
|  | SPROF-GO | <u>0.719</u> | <u>0.662</u> | 0.370 | <u>0.572</u> | <u>0.535</u> | 0.275 | <u>0.723</u> | <u>0.588</u> | 0.279 | <u>4.729</u> | <u>3.458</u> | 13.622 | 1.0 | 1.0 | 1.0 |
|  | TransFew | 0.696 | 0.627 | 0.398 | 0.537 | 0.514 | 0.334 | 0.616 | 0.516 | 0.273 | 5.186 | 3.878 | 13.881 | 1.0 | 1.0 | 1.0 |
| Single modal | Structure | 0.603 | 0.567 | 0.311 | 0.423 | 0.483 | 0.238 | 0.547 | 0.427 | 0.197 | 6.146 | 4.082 | 14.687 | 0.82 | 0.77 | 0.75 |
|  | Interpro | 0.579 | 0.592 | 0.435 | 0.434 | 0.499 | 0.384 | 0.546 | 0.459 | 0.300 | 6.219 | 3.932 | 12.620 | 0.80 | 0.86 | 0.79 |
|  | Sequence | 0.699 | 0.604 | 0.436 | 0.542 | 0.474 | 0.365 | 0.518 | 0.434 | 0.320 | 5.250 | 4.026 | 13.177 | 1.0 | 1.0 | 1.0 |
|  | Text | 0.663 | 0.623 | <u>0.502</u> | 0.536 | 0.511 | <u>0.450</u> | 0.638 | 0.517 | <u>0.397</u> | 5.110 | 3.597 | <u>11.561</u> | 0.86 | 0.90 | 0.90 |
|  | FunBind | **0.731** | **0.687** | **0.522** | **0.585** | **0.588** | **0.470** | **0.770** | **0.606** | **0.443** | **4.398** | **3.383** | **11.303** | 1.0 | 1.0 | 1.0 |

**Note:** FunBind: multimodal consensus with sequence, structure, text and interpro modality

**Fig. 5:**
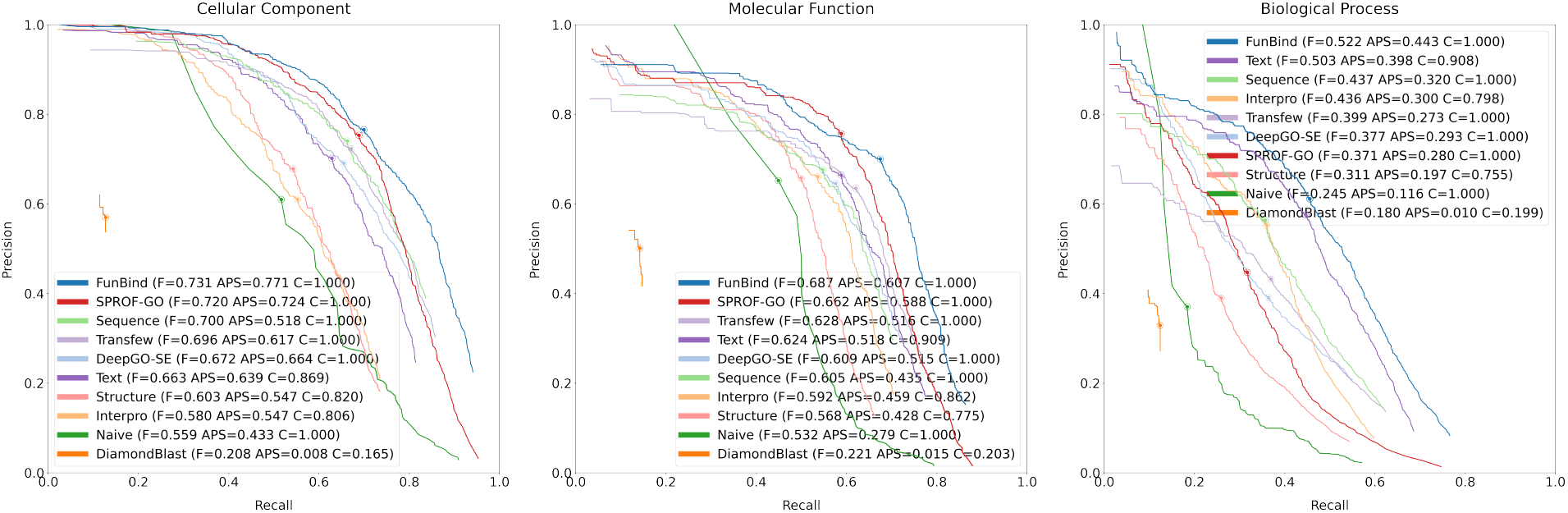
The Precision-Recall curves of FunBind and existing methods for the three ontologies (BP, MF, and CC) on the Test Novel dataset. The circled dot marks the point where each method achieves the highest *F*_*max*_. Each plot also includes the AUPR score and the coverage of each method. FunBind outperform all existing methods, particularly leading in BP by a large margin.

Like on Test All dataset, FunBind outperforms all the methods for the three function categories in terms of all the metrics. Particularly, it performs better than all baseline and current state-of-theart methods for BP by a large margin. It performs slightly better than Text for BP, while it substantially outperforms Text for both MF and CC. This confirms that the major improvement gained by FunFund for BP comes from Text, while combining Text with other modalities substantially improves its performance for MF and CC. Moreover, the performance of FunBind on Test Novel is similar to that on Test All, indicating that it is rather robust and generalizes well to novel proteins that are very dissimilar to the proteins in the training data.

## Discussion

Our results demonstrate that the multimodal model combining sequence, structure, textual descriptions, and InterPro domain annotations can enable accurate unsupervised zero-shot protein function prediction across all three Gene Ontology (GO) categories and further improve supervised protein function classification over the current state-of-the-art methods. The analysis reveals distinct contributions from individual modalities and their complementarity. Particularly, Text modality makes outstanding contributions to Biological Process (BP) and Molecular Function (MF) and valuable contributions to Cellular Component (CC), while Sequence and InterPro annotations can make stable contributions to all function categories as well. Structure, in its current form (3Di string), contributes modestly, but still can help improve the overall prediction accuracy when combined with other modalities. Overall, the four modalities capture some unique information about protein function and combining them generally works better than using one single modality.

Despite these promising findings, several avenues exist for future improvement. First, rather than relying on frozen encoders for initial representation generation, fine-tuning the base encoders, such as ESM for sequences, ProtsT5 for structure, and LLaMA for text, may further improve performance by adapting them to the function prediction task. Even selectively finetuning some of them might offer some gains without the full cost of joint training.

Second, the structural modality may be underutilized in our current implementation. The 3Di representation may be too coarse to offer higher prediction power than other modalities. A more detailed representation of protein structures, such as graph-based representations and atomic-level embeddings, may provide more robust structural signals that contribute more to function prediction. A more advanced structure encoder that directly generates structural embeddings from a 3D protein structure may further enhance function prediction.

Third, the multimodal foundational model could be extended to incorporate additional modalities that capture orthogonal biological information. One promising direction is the integration of protein–protein interaction (PPI) networks[22], which can encode relational context and functional associations often missed by individual protein-level descriptors. Combining PPI data with sequence, structure, domain annotations and text could unlock new capabilities, especially for inferring biological processes that depend on protein-protein interactions a lot.

Finally, in addition to protein function prediction, we envision FunBind can be used to generate the unique, unified representations of protein sequences, structures, text description, and Interpro domain annotations with protein function contexts for various downstream protein bioinformatics tasks, such as protein interaction prediction and protein design.

## Methods

### Datasets

We collected protein sequences and their functional annotations from the fifth Critical Assessment of Functional Annotation (CAFA5) [23] dataset, which primarily includes annotations released by November 2022, for model training and validation. Only sequences associated with at least one additional modality were retained for these purposes.

The predicted structures for the proteins were retrieved from AlphaFoldDB [24, 25], specifically the swissprot pdb v4.tar subset, and converted into one-dimensional structural representations — known as 3Di strings (sequences) — using FoldSeek [26–28]. These 3Di strings were used to generate structural embeddings.

Textual protein descriptions were sourced from UniProt/Swiss-Prot[29, 30], focusing on curated fields such as subcellular location, subunits, functions, induction, tissue specificity, and similarity. Additionally, InterPro domain annotations were extracted from the InterPro database[31] for proteins with available information. The statistics for the training data is presented in **Table 9**.

**Table 9:** The Statistics of Training Data.

| Ontology | Sequence | Structure | Text | InterPro |
| --- | --- | --- | --- | --- |
| CC | 92912 | 85491 | 75625 | 85722 |
| MF | 78637 | 72406 | 68500 | 74422 |
| BP | 92210 | 80243 | 74519 | 82370 |
| Pretraining | 133339 | 125021 | 110425 | 126861 |

To construct the test datasets, we collected newly annotated proteins from UniProt (specifically the goa uniprot all.gaf set), released on December 21, 2024. The GO terms were obtained from UniProt, while the Gene Ontology graph was sourced from the Gene Ontology Resource [8, 9]. To compile the complete set of GO terms (labels) for each protein, we first retrieved its direct annotations from UniProt and then expanded this set by traversing the GO graph to include all ancestor terms.

Following the guidelines established in CAFA [23], we retained only annotations supported by strong evidence codes (EXP, IDA, IPI, IMP, IGI, IEP, TAS, IC, HTP, HDA, HMP, HGI, HEP), ensuring that each protein is labeled with highconfidence functional information. We refer to this curated dataset of newly annotated proteins as **Test All** (see its statistics in **Table 5**). Moreover, the proteins in Test All that have less than 30% sequence identity with any proteins in the training data were used to create a dataset **Test Novel**. Test All and Test Novel were used to compare the fined-tuned FunBind and existing methods.

To evaluate zero-shot function prediction, we constructed another test set **(Test Zero)** from Test All. We collected all proteins that have at least one novel GO term that does not appear in the training data. This ensures that the evaluation measures the model’s ability to generalize to novel GO terms never encountered during training. The Test Zero data includes 659 proteins. Among them, 450 proteins have one new annotation (GO term), 156 have two, 21 have three, 24 have four, and 8 have five to six new annotations. Notably, protein A8BPK8 has 17 new annotations. The statistics of Test Zero is shown in **Table 1**, which details the number of proteins in three functional categories: CC (Cellular Component), MF (Molecular Function), and BP (Biological Process) and the All function categories combined.

### Modality Encoding

The overall architecture of FundBind is illustrated in **Figure 1**. It begins with an input modality encoding block, which generates initial embeddings for each protein by applying a modality-specific encoder to every available modality.

For protein sequences, we tested two pretrained protein language models: ESM2-t48 and ProstT5 [33]. While ProstT5 demonstrated superior performance in binding sequence embeddings to structural embeddings, ESM2-t48 achieved stronger results when aligning sequence embeddings with text and InterPro modalities, as shown in supplementary **Figure S8**. Based on the overall performance across all modalities, ESM2t48 was chosen as the default sequence encoder for the final model.

For protein structures, we employed FoldSeek [26–28, 34] to convert three-dimensional structures into 1D structural representations known as 3Di strings consisting of 3Di-tokens. These structural sequences were then encoded using the ProstT5 structural module, producing embeddings enriched with structural context.

Inspired by ProtST [35], we constructed textual descriptions by aggregating information from diverse protein-related fields, including protein names, functional descriptions, subcellular locations, subunits, induction, tissue specificity, and similarity. These fields were concatenated into a single textual sequence for each protein. The statistics of the fields is shown in **Table 10**. Supplementary **Figure S9** presents an example with all five modalities including textual descriptions.

**Table 10:** Summary of textual descriptions on the training Data. 110425 proteins in the training data have some textual information.

|  | Text Field |  |  |  |  |  |  |  |
| --- | --- | --- | --- | --- | --- | --- | --- | --- |
|  | Name | Desc. | Func. | Subunit | Sub. Loc. | Induc. | Tiss. Spec. | Sim. |
| # of proteins | 110425 | 110425 | 81618 | 53990 | 90246 | 12773 | 31163 | 88831 |
| Coverage | 100% | 100% | 73.9% | 48.9% | 81.7% | 11.6% | 28.2% | 80.4% |

Similarly, each protein’s InterPro annotation in text format was generated by combining the name, description, and the set of mapped InterPro domains, as illustrated in supplementary **Figure S9**.

For GO terms, we collected their Gene Ontology (GO) definitions spanning the three function categories: Cellular Component (CC), Molecular Function (MF), and Biological Process (BP). For each protein, we sampled a subset of GO terms preferably leaf nodes in the GO graph, and constructed textual representations by concatenating their namespaces, GO term names, and definitions, as depicted in supplementary **Figure S9**.

For both InterPro annotations and textual descriptions, we compared LLaMA2 [36], a large autoregressive language model, with BioBERT [37], a transformer model pre-trained specifically on biomedical literature. Based on its superior performance across modalities shown in supplementary **Figure S8**, we selected LLaMA2 as the encoder for the two modalities. Similarly, for generating GO term embeddings, we used LLaMA2 to encode the textual descriptions of GO terms.

When using LLaMA2, each input annotation is tokenized with an End-of-Sequence (EOS) token appended to mark the end, and sequences are padded to ensure consistent batching. We set the max length parameter to 1024 tokens for all three modalities, as most input sequences fall within this range. The distribution of sequence lengths is shown in supplementary **Figure S10**.

To reduce computational costs, we kept the weights of all base encoders (ESM, ProstT5, and LLaMA2) frozen throughout the training of FunBind. This design choice allows it to focus on learning effective cross-modality alignments and function prediction.

### Modality Alignment (Binding)

The modality alignment and integration module aims to learn a unified embedding space that facilitates cross-modality alignment. We employ a pairwise contrastive learning approach, using pairs of modalities (S, M), where S represents protein sequences and M represents a complementary modality. In our experiments, we bind structural, textual, InterPro domain annotations, and GO terms modalities to the central sequence modality. The objective is to maximize the similarity between two paired modalities from the same proteins in the embedding space while minimizing the similarity between two unpaired (negative) modalities from different proteins, thereby encouraging modality embeddings to align when semantically related.

Given a protein *i*, with sequence *S*_*i*_ and another modality *M*_*i*_, we generate embeddings as follows: *q*_*i*_ = *f* (*S*_*i*_) and *k*_*i*_ = *g*(*M*_*i*_), where *f* and *g* are modality-specific projection networks applied to the representations obtained during the encoding phase using pre-trained encoders (i.e., language models). These projection layers transform the modality-specific embeddings into a shared embedding space suitable for contrastive learning.

To capture modality-specific nuances and enable effective cross-modality integration, we employ a Mixture of Experts (MoE) architecture for each input modality, as illustrated in **Figure 6**. Each MoE module comprises 1 to 3 parallel experts, with each expert implemented as a stack of multilayer perceptrons (MLPs) interleaved with Batch Normalization, GELU (Gaussian Error Linear Unit) activations, and Dropout layers.

**Fig. 6:**
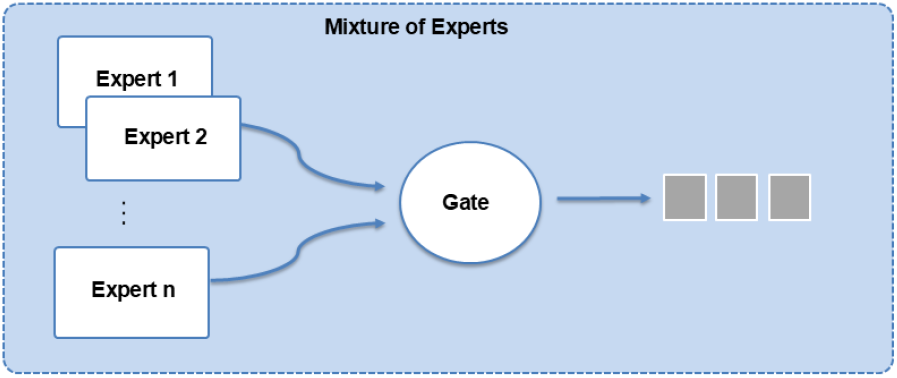
Illustration of the Mixture of Experts (MoE) architecture. Multiple expert networks process the input in parallel, and a gating mechanism dynamically weights their outputs to produce the final representation.

The outputs of all experts are combined using a gating mechanism, which dynamically computes a weighted combination of the expert outputs based on the input. This gating network assigns higher weights to experts that are more relevant for a given input, enabling the model to flexibly adapt to different data patterns within each modality. As a result, each expert can specialize in capturing distinct features, while the gating mechanism ensures that the final representation is tailored to the specific characteristics of the input (as illustrated in **Figure 1**).

To promote the learning of complementary representations and prevent experts from collapsing to similar functions, we introduce a diversity loss. This loss encourages the gating network to activate experts in a way that maximizes variation among their outputs. It is defined as:

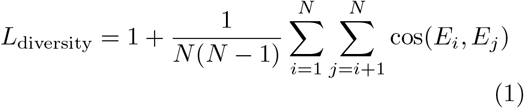

where *E*_*i*_ and *E*_*j*_ are the output embeddings of experts *i* and *j*, and *cos*(*E*_*i*_, *E*_*j*_) denotes their cosine similarity.

This loss penalizes similarity between expert outputs—growing larger when embeddings are highly similar and smaller when they are dissimilar or negatively correlated. It encourages each expert to specialize in different aspects of the input, thereby enriching the model’s ability to capture diverse, modality-specific representations.

### Self-supervised Pretraining

During self-supervised pretraining, we optimize the embeddings generated by a MoE using an InfoNCE loss for contrastive learning [12, 34, 38], regularized by the diversity loss term to ensure the experts produce diverse representations. The InfoNCE loss(*L*_*S,M*_) is defined as:

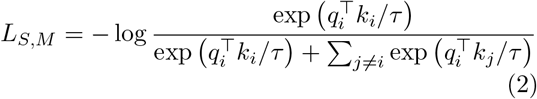

where *τ* is a scalar temperature controlling the smoothness of the softmax, *q*_*i*_ represents the embedding of protein *i* from the sequence modality. *k*_*i*_ represents the embedding of protein *i* from non-sequence modality *M, k*_*j*_ represents the embedding of protein *j* (where *j≠ i*) from modality *M* serving as a negative sample.

The final loss function, *L*_*total*_, is a combination of the InfoNCE loss and the diversity loss:

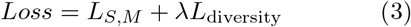

*Loss* = *L*_*S,M*_ + *λL*_diversity_ (3) where *λ* is a hyperparameter that controls the weight of the diversity loss term.

### Unsupervised Zero-Shot Function Prediction

After FunBind is pretrained, it can be directly used to make zero-shot prediction of protein function terms from any non-function modality (e.g., sequence and text) by calculating the similarity between the embeddings generated for candidate GO terms and the embeddings of the other modality of proteins. The similarity can be used to rank GO terms for a protein. Top *k* GO terms can be used as predictions, no matter if they were encountered during the training or not. Therefore, FunBind can predict novel GO terms never seen before.

### Supervised Function Classification via Fine-Tuning

The pretrained FundBind can also be fine-tuned to predict protein function (GO terms) from one or multiple available non-function modalities. Building on the pretrained multimodal backbone, we extend the model to perform supervised multilabel function classification for the three GO function categories: CC, MF, and BP. Different from the open-ended unsupervised zero-shot prediction that works for any GO terms including novel ones, the supervised classification requires the number of function classes (terms) is predetermined. Like existing supervised function prediction methods, to mitigate label sparsity and focus on informative GO terms, we limit the prediction space to the frequently annotated GO terms in the training data. Specifically, we retain GO terms with at least 20 annotations for CC and MF, and a higher threshold of 250 annotations for BP, resulting in 1,043 classes for CC, 1,529 classes for MF, and 1,631 classes for BP.

To support this supervised learning task, we augment the pretrained foundational model backbone with a dedicated classification head for each non-function modality. Each classification head is composed of a series of multilayer perceptrons (MLPs) interleaved with Batch Normalization, GELU activation functions, and Dropout layers.

The final layer outputs logits for the GO terms for each specific ontology (CC, MF or BP) and applies a sigmoid activation to enable multi-label prediction of GO terms. Because proteins can have a varied number of input modalities, the model generates one separate prediction for each available non-function modality. These classification heads allow the general-purpose features learned during the pretraining to be specialized for the gene ontology-specific function prediction.

We jointly optimize all classification heads using the binary cross-entropy (BCE) loss, formulated as:

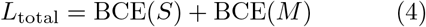

where: BCE(*S*) denotes the binary cross-entropy loss for the sequence modality, and BCE(*M*) refers to the loss computed for another available modality *M*, such as structure, text, or InterPro. At inference time, the model supports flexible prediction using any single modality or a combination of available modalities. This design allows it to handle varying levels of data completeness and leverage multiple sources of information when available for more robust predictions. While there are many possible strategies for combining the predictions of individual modalities, in this work, we adopt a simple weighted averaging scheme to combine individual predictions. Specifically, we assign weights ranging from 0.0 to 1.0 in increments of 0.1 to the predictions of individual modalities and ensure the sum of the weights equals to 1.

We obtained the optimal weight configurations using the validation data by evaluating all valid combinations and selecting the one that achieves the highest weighted F-measure when all modalities are used (FunBind). The optimal combination weights for FunBind are reported in supplementary **Table S1**. The weights were blindly applied to Test All and Test Novel datasets to obtain the test results to evaluate their performance.

## Data Availability

The original data used in this study can be downloaded using the script available in our https://github.com/jianlin-cheng/FunBind. The repository also includes scripts for generating the training and test datasets from the original data.

## Code Availability

The source code for FunBind is available at the GitHub repository: https://github.com/jianlincheng/FunBind.

## Acknowledgments

This work was supported in part by U.S.A. NSF grants (grant #: DBI2308699 and CCF2343612).

## Author Contributions Statement

JC conceived the project. JC and FB designed the method. FB implemented the method, performed the experiment, and collected the data. FB, JC, and YW analyzed the data. FB and JC wrote the manuscript.

