## Supplementary Material for "A unified multimodal model for generalizable zero-shot and supervised protein function prediction"

FUNBIND

–

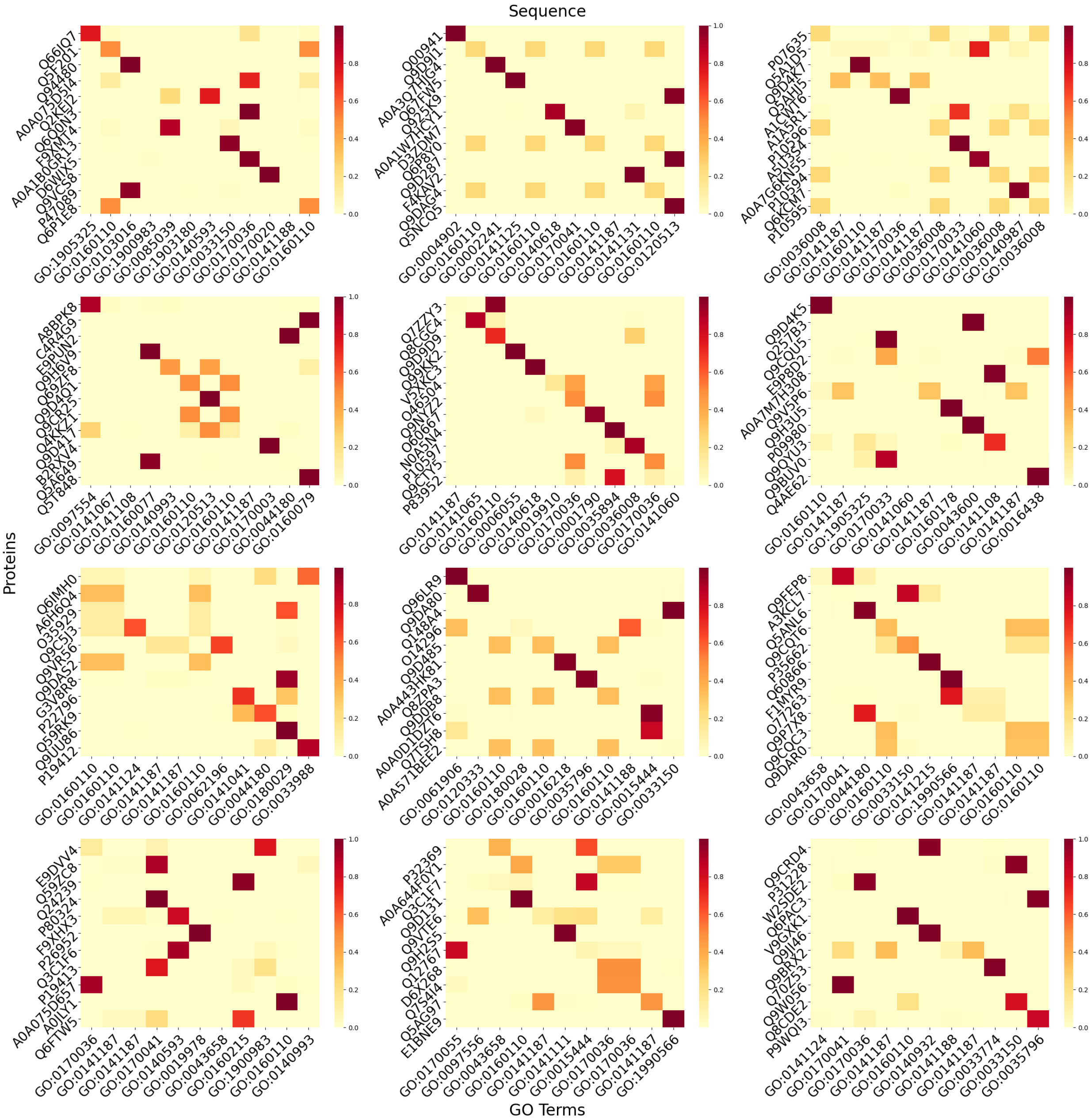


Figure S1: Heatmaps illustrating the similarity between proteins and GO terms for 12 groups, using the Sequence modality for retrieval. In each heatmap, rows correspond to proteins and columns to GO terms. The diagonal elements indicate the similarity between each protein and its corresponding true GO term.

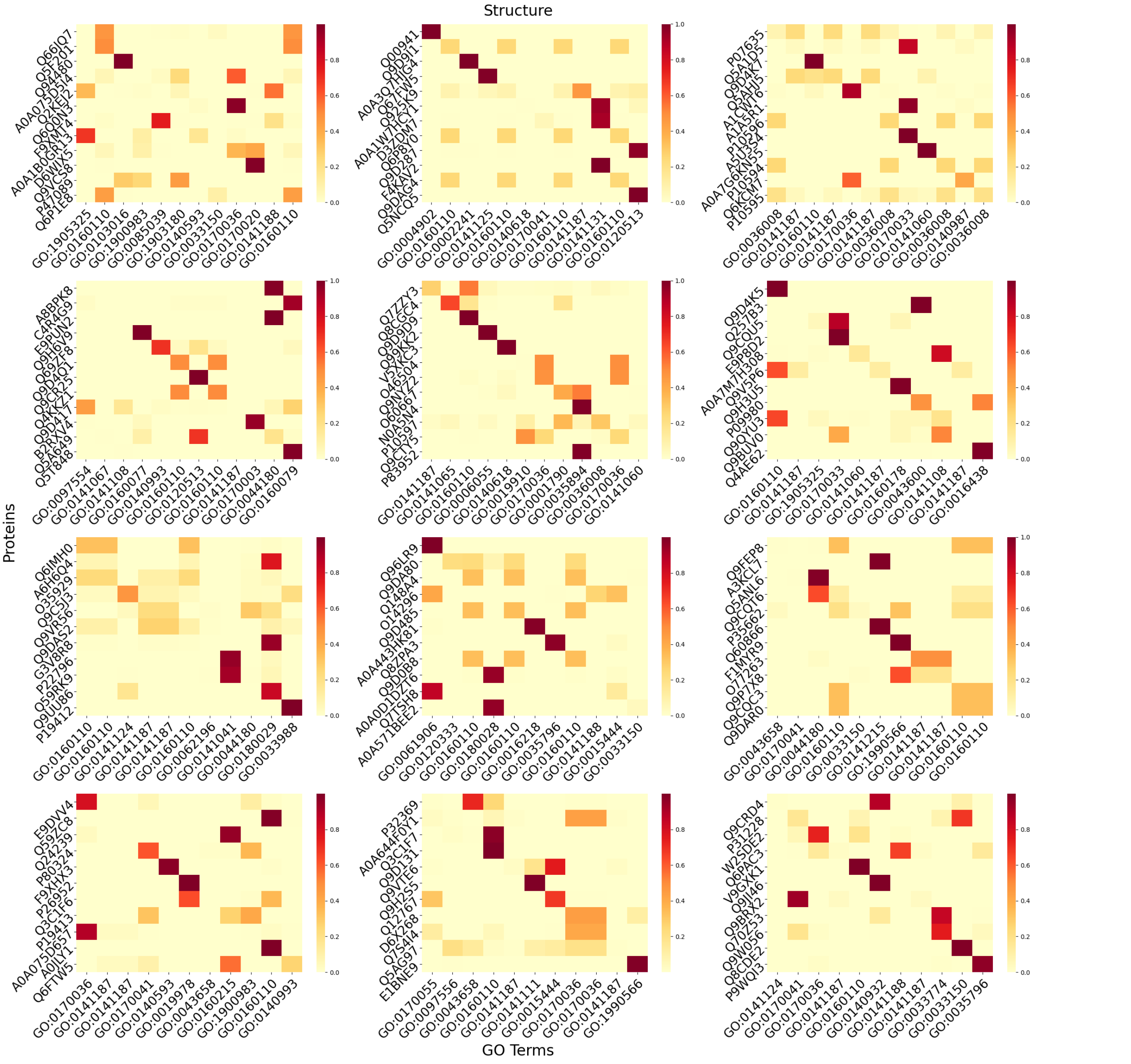


Figure S2: Heatmaps illustrating the similarity between proteins and GO terms for 12 groups, using Structure modality for retrieval. Each heatmap presents a similarity matrix, where rows correspond to proteins and columns to GO terms. The diagonal elements indicate the similarity between each protein and its true GO term.

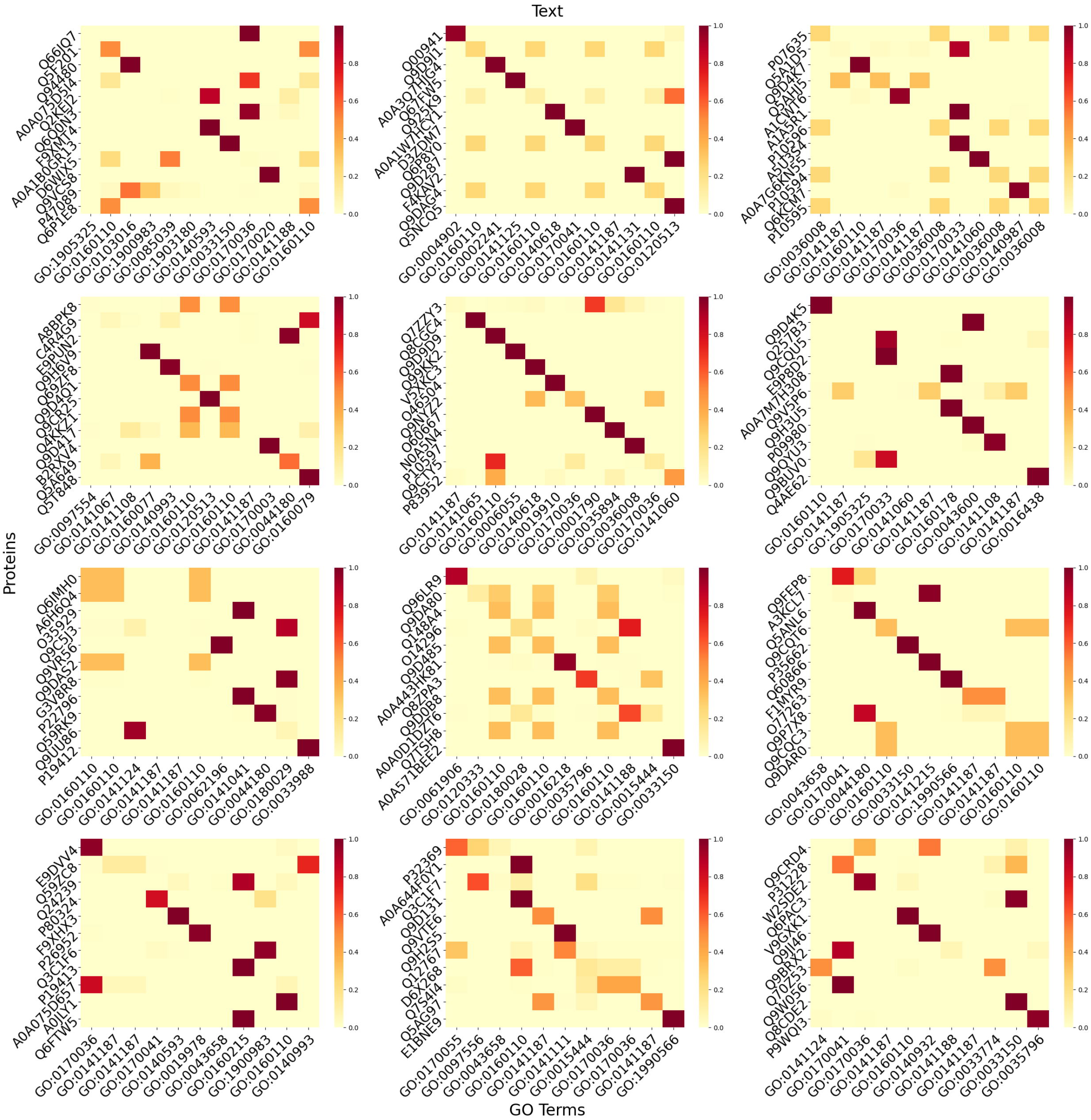


Figure S3: Heatmaps illustrating the similarity between proteins and GO terms for 12 groups, using Text modality for retrieval. Each heatmap presents a similarity matrix, where rows correspond to proteins and columns to GO terms. The diagonal elements indicate the similarity between each protein and its true GO term.

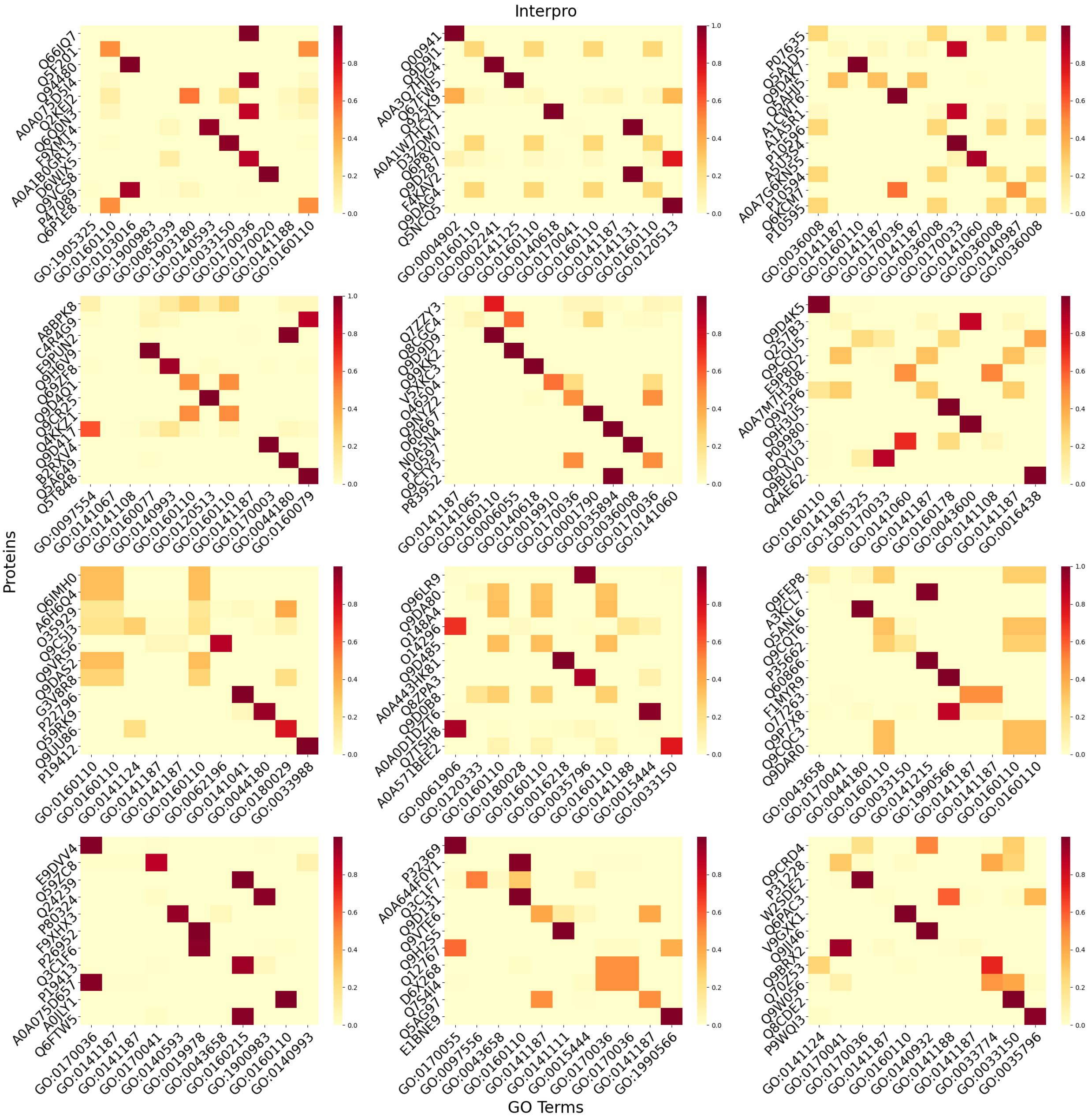


Figure S4: Heatmaps illustrating the similarity between proteins and GO terms for 12 groups, using Interpro modality for retrieval. Each heatmap presents a similarity matrix, where rows correspond to proteins and columns to GO terms. The diagonal elements indicate the similarity between each protein and its true GO term.

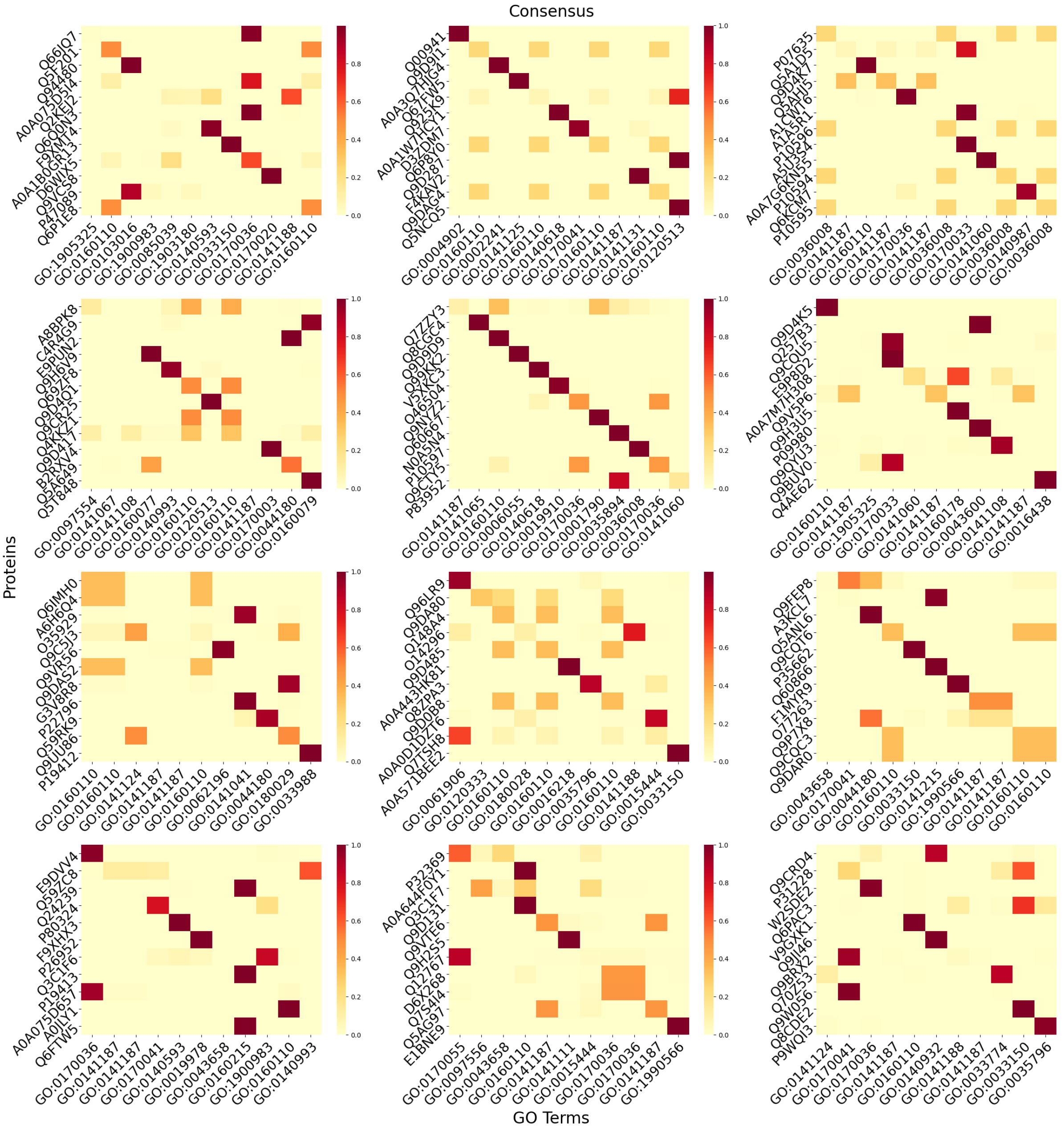


Figure S5: Heatmaps illustrating the similarity between proteins and GO terms for 12 groups, using the consensus predictions of all four modalities for retrieval. Each heatmap presents a similarity matrix, where rows correspond to proteins and columns to GO terms. The diagonal elements indicate the similarity between each protein and its true GO term.

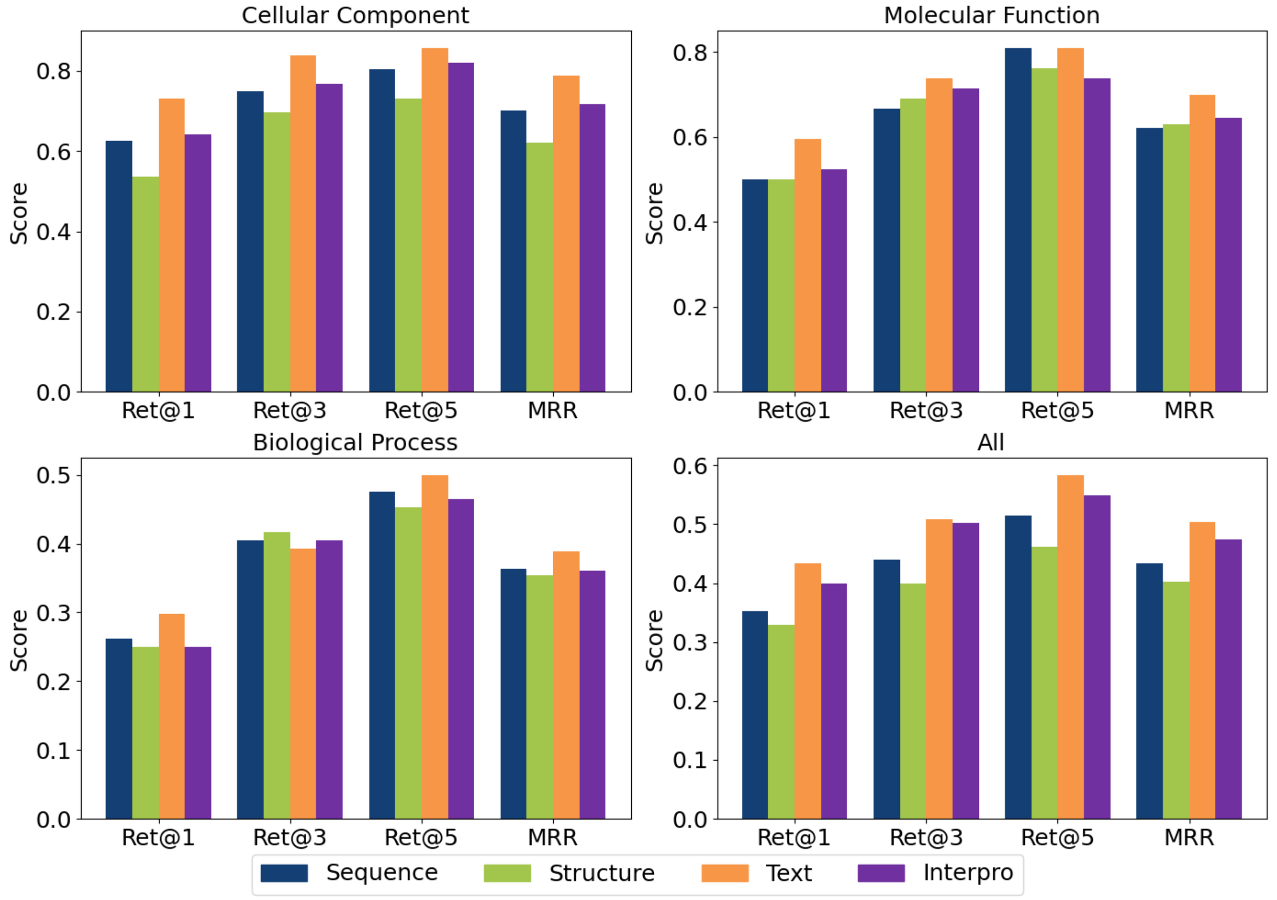


Figure S6: Zero-shot prediction performance for the test proteins with all modalities available in Test_zero dataset for each of the three GO function categories (Cellular Component, Molecular Function, Biological Process) and All function categories combined using individual modalities as queries. FunBind predict GO terms for all proteins in a single batch without dividing them into groups. The performance is reported using Ret@1, Ret@3, Ret@5, and MRR metrics. Each color represents a different modality: sequence, structure, text, and InterPro domain annotations.

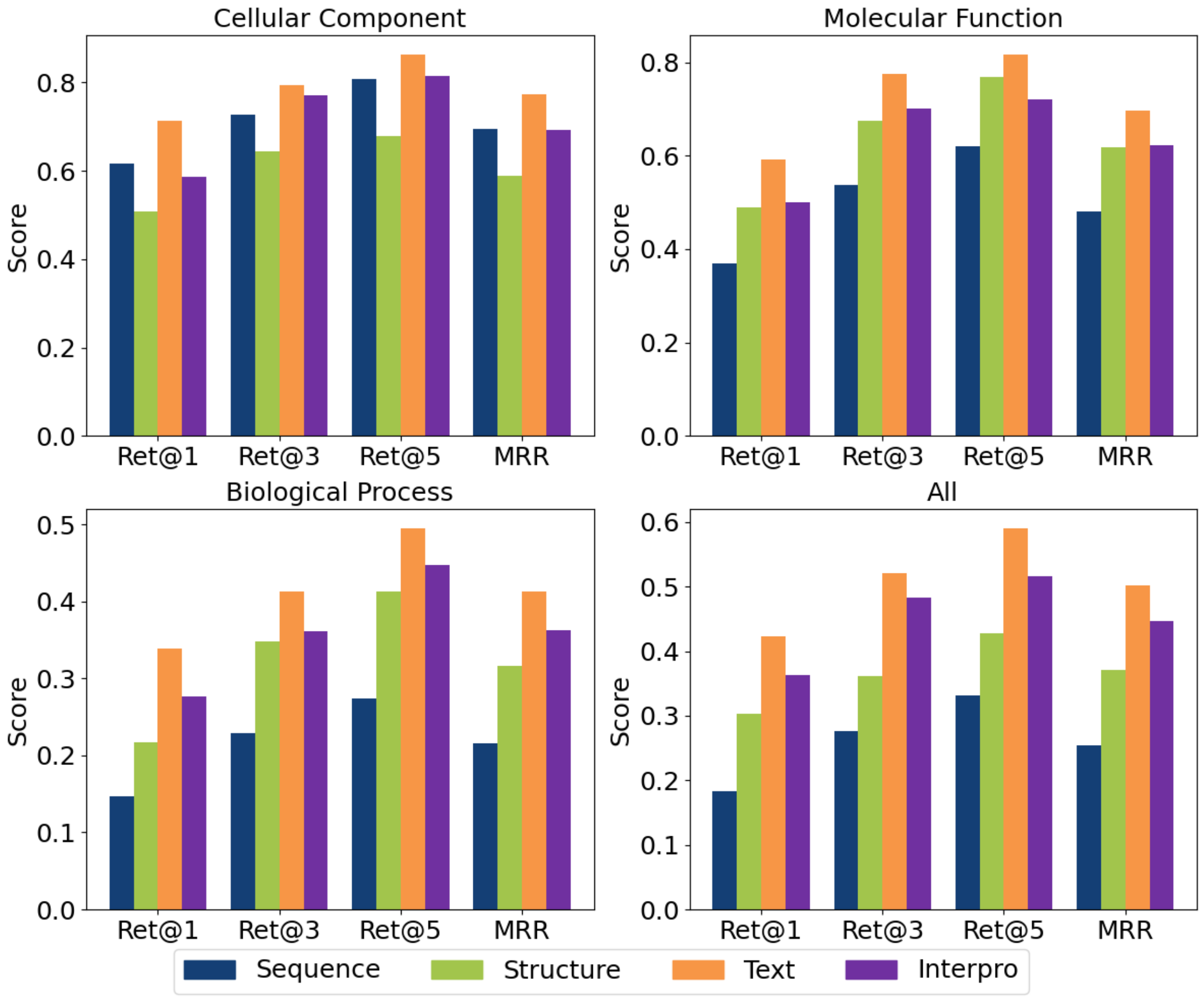


Figure S7: Zero-shot prediction performance on the entire test proteins((irrespective of modality availability) in Test_Zero dataset for each of the three GO function categories (Cellular Component, Molecular Function, Biological Process) and All function categories combined using individual modalities as queries. FunBind predict GO terms for all proteins in a single batch without dividing them into groups. The performance is reported using Ret@1, Ret@3, Ret@5, and MRR metrics.

### Supplementary Note 1: Evaluation Metrics

In this work, we evaluate protein function prediction and zero-shot retrieval using three CAFA metrics—$F_{\max}$, $S_{\min}$, and weighted $F_{\max}$ [3, 7]. Additionally, we report the area under the precision-recall curve (AUPR) and standard retrieval metrics, including Recall@$k$ (Ret@$k$) and Mean Reciprocal Rank (MRR). These metrics are defined as follows.

- **Precision**

$$\mathrm{pr}(\tau) = \frac{1}{m(\tau)} \sum_{i=1}^{m(\tau)} \frac{\sum_f \mathbb{I}(f \in P_i(\tau) \wedge f \in T_i)}{\sum_f \mathbb{I}(f \in P_i(\tau))}$$

- **Recall**

$$\mathrm{rc}(\tau) = \frac{1}{n_e} \sum_{i=1}^{n_e} \frac{\sum_f \mathbb{I}(f \in P_i(\tau) \wedge f \in T_i)}{\sum_f \mathbb{I}(f \in T_i)}$$

- **$\mathbf{F_1}$ Score**

$$F_{1(\tau)} = 2 \times \frac{\mathrm{pr}(\tau) \times \mathrm{rc}(\tau)}{\mathrm{pr}(\tau) + \mathrm{rc}(\tau)}$$

- **Maximum $\mathbf{F_1}$ Score**

$$F_{\max} = \max_{\tau} (F_1(\tau))$$

where $f$ is a term, $P_i(\tau)$ is the set of predictions, $T_i$ denotes the corresponding ground-truth, $i$ represents the protein sequence under consideration, and $\tau$ is the decision threshold. $m(\tau)$ is the number of proteins sequences with at least one predicted score greater than or equal to the decision threshold $\tau$, $\mathbb{I}(\cdot)$ is an indicator function, and $n_e$ is the number of proteins in the test set for a particular test study.

- **Information Content** (*ic*) of term $f$ is computed as:

$$\mathrm{IC}(f) = \log_2 \frac{1}{\Pr(f|P(f))}$$

- **Weighted precision**:

$$\mathrm{wpr}(\tau) = \frac{1}{m(\tau)} \sum_{i=1}^{m(\tau)} \frac{\sum_f ic(f) \cdot \mathbb{I}(f \in P_i(\tau) \wedge T_i(\tau))}{\sum_f ic(f) \cdot \mathbb{I}(f \in P_i(\tau))}$$

- **Weighted Recall**:

$$\mathrm{wrc}(\tau) = \frac{1}{n_e} \sum_{i=1}^{n_e} \frac{\sum_f ic(f) \cdot \mathbb{I}(f \in P_i(\tau) \wedge T_i(\tau))}{\sum_f ic(f) \cdot \mathbb{I}(f \in T_i(\tau))}$$

Here, $\Pr(f|P(f))$ represents the probability that term $f$ in the ontology is associated with a protein given that all of its parents are associated.

- **Remaining Uncertainty**

$$ru(\tau) = \frac{1}{ne}\sum_{i=1}^{ne}\sum_{f} ic(f) \cdot \mathbb{I}(f \notin P_i(\tau) \wedge f \in Ti)$$

- **Missing Information**

$$mi(\tau) = \frac{1}{ne}\sum_{i=1}^{ne}\sum_{f} ic(f) \cdot \mathbb{I}(f \in P_i(\tau) \wedge f \notin Ti)$$

- $S_{min}$

$$S_{min} = \min_{\tau} \sqrt{ru(\tau)^2 + mi(\tau)^2}, \tau$$

- **Area under precision recall curve (AUPR)**

$$\text{AUPR} = \int_0^1 \text{Precision}(R)\, dR$$

where Precision(R) represents the precision at a given recall level (R).

- **Recall@k**

$$\text{Recall@}k = \frac{1}{N}\sum_{i=1}^{N} \mathbf{1}\left[\exists j \in \mathcal{G}_i : j \in \text{topk}(s_i, k)\right]$$

where $N$ is the number of queries (batch size), $s_i$ is the similarity scores for the $i$-th query against all candidates, $\mathcal{G}_i$ is the set of ground truth indices for the $i$-th query, $\text{topk}(s_i, k)$ returns the indices of the $k$ highest scoring candidates, and $\mathbf{1}[\cdot]$ is the indicator function that returns 1 if the condition is true, 0 otherwise.

- **Mean Reciprocal Rank**

$$\text{MRR} = \frac{1}{N}\sum_{i=1}^{N} \frac{1}{\min_{j \in \mathcal{G}_i} \text{rank}(j, s_i)}$$

where $N$ is the number of queries, $\mathcal{G}_i$ is the set of ground truth indices for the $i$-th query, $\text{rank}(j, s_i)$ returns the position of candidate $j$ in the sorted (descending) list of similarity scores $s_i$, and $\min_{j \in \mathcal{G}_i}$ finds the minimum rank among all ground truth indices.

#### Supplementary Note 2: Existing Methods

For full training, we compared our method to five baseline methods, namely Naive, DiamondBLAST [2, 5], TransFew [1], DeepGO-SE [4], and SPROF-GO [6]. Here's a concise overview of each method:

Naive: The Naive method simply uses the frequency of Gene Ontology (GO) terms in the training dataset to make predictions.

DiamondBLAST [2, 5] is based on sequence similarity scores obtained through BLAST, it identifies similar sequences from the training set and transfers annotations from the most similar ones .

SPROF-GO [6] is an alignment-free method employing a pre-trained protein language model to extract informative sequence embeddings. It utilizes self-attention pooling to focus on crucial residues and integrates homology information using a label diffusion algorithm . Test predictions were obtained through the provided web server for SPROF-GO.

DeepGO-SE [4] utilizes a pre-trained large protein language model combined with GO background knowledge and protein-protein interactions (PPIs) to make accurate predictions about protein functions. Predictions for DeepGO-SE were generated by cloning and running the tool locally.

TransFew [1] is a deep learning method for protein function prediction, with a focus on rare GO terms. It generates protein and GO term embeddings using pretrained models (ESM2 and BioBERT), and integrates them via cross-attention to transfer knowledge from common to rare terms and improve prediction accuracy.

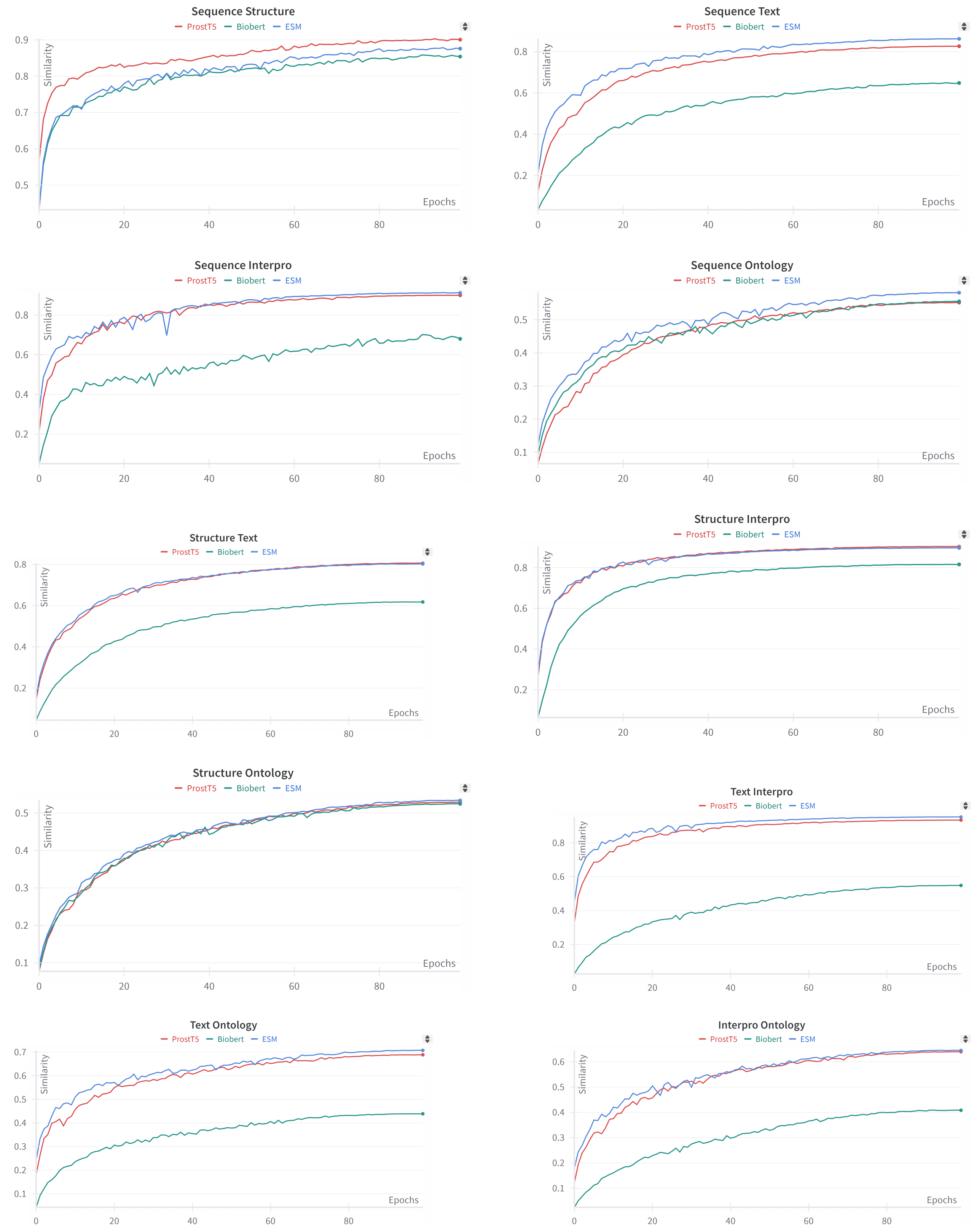


Figure S8: Cross-modal similarity on the validation set during pretraining, illustrating the alignment between modality embeddings across epochs. Each plot shows the cosine similarity between pairs of modality embeddings. When ”ProstT5” indicates that ProstT5 was used as the sequence encoder; similarly, ”BioBERT” indicates the use of BioBERT as the base encoder for Text and Interpro. In all other cases, the default encoders are: ESM for sequence, ProstT5 for structure, and LLaMA2 for text, InterPro, and ontology.

**Example Protein -- Q9UJZ1**

**Sequence**: MLARAARGTGALLLRGSLLASGRAPRRASSGLPRNTVVLFVPQQEAWVVERMGR **...** KMS
**Structure:** dddddddddddddddddddddddpddppppfdaqdqkdaaapqkwwfkdfpnh **...** pvd
**Textual Annotation:** **PROTEIN NAME**: STML2_HUMAN; **DESCRIPTION**: Stomatin-like protein 2, mitochondrial; **FUNCTION**: Mitochondrial protein that probably regulates the biogenesis and the activity of mitochondria. Stimulates cardiolipin biosynthesis, binds cardiolipin-enriched membranes where it recruits and stabilizes some proteins including prohibitin and may therefore act in the organization of functional microdomains in mitochondrial membranes. Through regulation of the mitochondrial function may play a role into several biological processes including cell migration, cell proliferation, T-cell activation, calcium homeostasis and cellular response to stress. May play a role in calcium homeostasis through negative regulation of calcium efflux from mitochondria. Required for mitochondrial hyperfusion a pro-survival cellular response to stress which results in increased ATP production by mitochondria. May also regulate the organization of functional domains at the plasma membrane and play a role in T-cell activation through association with the T- cell receptor signaling complex and its regulation.; **SUBUNIT**: Forms homooligomers. Interacts with MFN2; may form heterooligomers. Interacts with CACNA2D2 (By similarity). Interacts with PHB1 and PHB2; recruits them to cardiolipin-enriched mitochondrial membranes and stabilizes them.; **SUBCELLULAR LOCATION**: Cell membrane ; Peripheral membrane protein . Mitochondrion . Mitochondrion inner membrane ; Lipid-anchor . Mitochondrion intermembrane space . Membrane raft . Cytoplasm, cytoskeleton . Note=Behaves as an integral membrane protein of the mitochondrion despite the absence of a detectable transmembrane domain . Also associates with the actin cytoskeleton and membrane rafts in activated T-cells (PubMed:10713127, PubMed:18641330). A minor pool is associated with the plasma membrane and is enriched at the immunological synapse in activated T-cells.; **INDUCTION**: Up-regulated in activated B- and T-cells and upon mitochondrial stress by chloramphenicol.; **TISSUE SPECIFICITY**: Ubiquitously expressed at low levels. Expressed in lymphoid tissues (at protein level).; **SIMILARITY**: Belongs to the band 7/mec-2 family.
**Interpro:** **Protein Name**: STML2_HUMAN; **Description**: Stomatin-like protein 2, mitochondrial; **Interpro Name**: Band 7/SPFH domain superfamily, **Interpro Type**: Homologous_superfamily; **Interpro Name**: STML2-like, C-terminal extension, **Interpro Type**: Domain; **Interpro Name**: Band 7/mec-2 domain-containing protein, **Interpro Type**: Family; **Interpro Name**: Stomatin/HflK family, **Interpro Type**: Family; **Interpro Name**: Band 7 domain, **Interpro Type**: Domain
-------------------------------------------------------------------------------**zeroshot** -------------------------------------------------------------------------------
**Ontology:** **Cellular Component::** mitochondrial inner membrane: The inner, i.e. lumen-facing, lipid bilayer of the mitochondrial envelope. It is highly folded to form cristae. ; mitochondrial intermembrane space: The region between the inner and outer lipid bilayers of the mitochondrial envelope. ; actin cytoskeleton: The part of the cytoskeleton (the internal framework of a cell) composed of actin and associated proteins. Includes actin cytoskeleton-associated complexes. ; membrane raft: Any of the small (10-200 nm), heterogeneous, highly dynamic, sterol- and sphingolipid-enriched membrane domains that compartmentalize cellular processes. Small rafts can sometimes be stabilized to form larger platforms through protein-protein and protein-lipid interactions. ; immunological synapse: An area of close contact between a lymphocyte (T-, B-, or natural killer cell) and a target cell formed through the clustering of particular signaling and adhesion molecules and their associated membrane rafts on both the lymphocyte and the target cell and facilitating activation of the lymphocyte, transfer of membrane from the target cell to the lymphocyte, and in some situations killing of the target cell through release of secretory granules and/or death-pathway ligand-receptor interaction. **Molecular Funtion::** cardiolipin binding: Binding to cardiolipin. ; T cell receptor binding: Binding to a T cell receptor, the antigen-recognizing receptor on the surface of T cells. ; GTPase binding: Binding to a GTPase, any enzyme that catalyzes the hydrolysis of GTP. **Biological Process::** T cell receptor signaling pathway: The series of molecular signals initiated by the cross-linking of an antigen receptor on a T cell. ; mitochondrion organization: A process that is carried out at the cellular level which results in the assembly, arrangement of constituent parts, or disassembly of a mitochondrion; includes mitochondrial morphogenesis and distribution, and replication of the mitochondrial genome as well as synthesis of new mitochondrial components. ; intracellular calcium ion homeostasis: A homeostatic process involved in the maintenance of a steady state level of calcium ions within a cell. ; protein complex oligomerization: The process of creating protein oligomers, compounds composed of a small number, usually between three and ten, of component monomers; protein oligomers may be composed of different or identical monomers. Oligomers may be formed by the polymerization of a number of monomers or the depolymerization of a large protein polymer.

Figure S9: A protein example with five modalities: sequence, structure, text description, InterPro domain annotations, and Gene Ontology terms

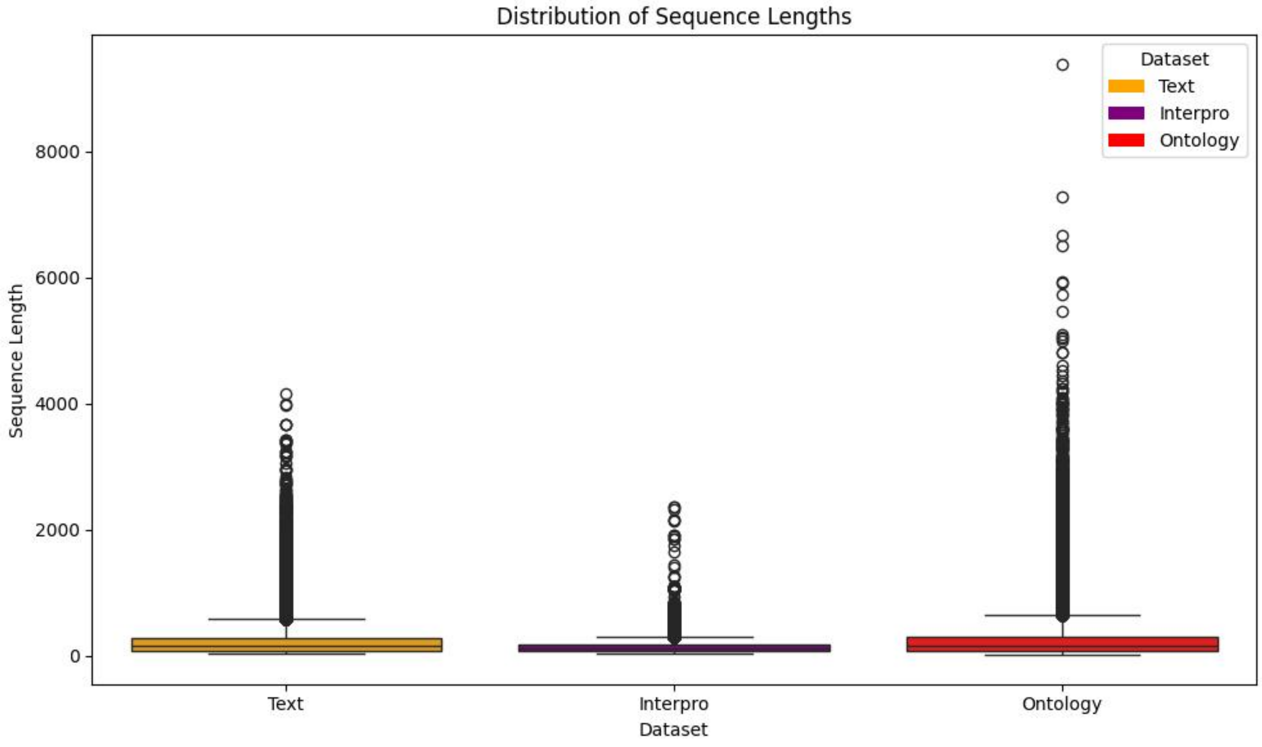


Figure S10: Distribution of text sequence lengths of three modalities: Text description, InterPro annotations, and Gene Ontology terms.

Table S1: The weights of the modalities for FunBind optimized on the validation dataset according to weighted F-measure.

| Method | Ontology | Sequence | Structure | Text | InterPro |
|---|---|---|---|---|---|
| FunBind | CC | 0.3 | 0.1 | 0.5 | 0.1 |
| | MF | 0.2 | 0.1 | 0.4 | 0.3 |
| | BP | 0.2 | 0.1 | 0.6 | 0.1 |

Table S2: Top modality weight configurations based on average Ret@1 across 10 runs.

| Ontology | Sequence | Structure | Text | InterPro |
|---|---|---|---|---|
| CC | 0.3 | 0.1 | 0.3 | 0.3 |
| MF | 0.1 | 0.1 | 0.5 | 0.3 |
| BP | 0.3 | 0.4 | 0.2 | 0.1 |
| All | 0.2 | 0.1 | 0.5 | 0.2 |
